# Distribution of metabotropic serotonergic receptors in GABAergic and glutamatergic neurons in the auditory midbrain

**DOI:** 10.1101/2025.09.14.675398

**Authors:** Karen L. Galindo, Zoya A. Nazir, Marina A. Silveira

## Abstract

The neurotransmitter serotonin modulates a variety of behavioral and physiological responses in the brain. Serotonergic neurons from the dorsal raphe nuclei send a dense projection to the auditory system, including the inferior colliculus (IC), the midbrain hub of the central auditory system. In the IC, serotonin alters how neurons respond to complex sounds, and it has been implicated in the generation or perception of tinnitus. However, the distribution of serotonergic receptors and the identity of neurons that express serotonergic receptors in the IC remains unclear. Here, we hypothesized that IC GABAergic and glutamatergic neurons differentially express serotonergic receptors. To test this hypothesis, we performed in situ hybridization in IC brain slices of male and female mice using probes for *Vgat* (GABAergic neuron marker) and *Vglut2* (glutamatergic neuron marker), along with probes for six subtypes of metabotropic serotonergic receptors: 5-HT1_A_ and 5-HT1_B_ (*Htr1a* and *Htr1b,* inhibitory, G_i/o_ G protein receptors), 5-HT2_A_, 5-HT2_B_ and 5-HT2_C_ (*Htr2a, Htr2b,* and *Htr2c* excitatory, G_q11_ G protein receptors) and, 5-HT7 (excitatory, G_s_ G protein receptors). Our data show that glutamatergic IC neurons primarily express inhibitory serotonergic receptors. In contrast, a larger proportion of GABAergic neurons express excitatory serotonergic receptors. Our data suggest that serotonin exerts an inhibitory net effect on IC neuronal circuits. These findings contribute to our understanding of how serotonin signaling influences auditory processing. The differential expression of serotonergic receptors may help shape the balance of excitation and inhibition in the auditory midbrain, affecting sound processing.

## Introduction

The serotonergic system influences many physiological processes, as well as pathological conditions such as anxiety, depression, obesity, autism, and Alzheimer’s disease (Meltzer et al., 1998; Oishi et al., 2010; Muller et al., 2016; van Galen et al., 2021; Tahiri et al., 2024). The majority of serotonergic neurons are concentrated in the dorsal raphe nuclei (DRN) and send projections to multiple brain regions, including the central auditory system (Thompson et al., 1994a). Serotonergic collaterals are found throughout the auditory pathway, and serotonergic signaling has significant impacts on neuronal responses in different auditory nuclei (Hurley et al., 2002; Tang and Trussell, 2015; Ko et al., 2016). In the auditory cortex, serotonin decreases the excitability of pyramidal neurons (Rao et al., 2010). In the dorsal cochlear nucleus, serotonin enhances the excitability of fusiform and vertical cells by activating 5-HT2_A/C_ and 5-HT7 receptors (Tang and Trussell, 2015, 2017). Serotonin also influences the initiation of action potentials in the medial superior olive (Ko et al., 2016). Serotonergic neurons send a major projection to the inferior colliculus (IC), the midbrain hub of the central auditory system (Klepper and Herbert, 1991; Hurley et al., 2002), which is comprised of GABAergic and glutamatergic neurons (Oliver et al., 1994; Ono et al., 2017). In the IC, serotonin binds to serotonergic receptors, resulting in either enhancement or suppression of auditory responses (Hurley, 2006; Bohorquez and Hurley, 2009; Ramsey et al., 2010). Additionally, serotonin differentially modulates responses to complex sounds, including vocalizations and direction selectivity for frequency-modulated sweeps (Hurley and Pollak, 1999; Hood and Hurley, 2023). A previous study showed that the concentration of extracellular serotonin in the IC increases rapidly after noise presentation and over the course of social interactions (Hall et al., 2010).

Dysfunction in serotonergic signaling is involved in tinnitus perception (Jarach et al., 2022) and generation (Stark et al., 1985; Shemen, 1998). Furthermore, medications that selectively inhibit the reuptake of serotonin (SSRIs), commonly used in the treatment of depressive disorders (Shemen, 1998), have been shown to improve tinnitus perception (Fornaro and Martino, 2010; Oishi et al., 2010). Additionally, activation of a specific subtype of serotonergic receptor has been shown to improve temporal processing in the auditory cortex of a mouse line with auditory hypersensitivity (Tao et al., 2025). However, despite the importance of serotonergic modulation in the IC, the distribution of serotonergic receptors and which subtypes of serotonergic receptors are expressed by GABAergic and glutamatergic IC neurons remain unknown.

The modulatory impact of serotonin across the brain is driven by the selective expression of serotonin receptor types on specific cell populations, which in turn influences the function of neural networks (Andrade, 1998; Santana et al., 2004; Winterer et al., 2011). For example, serotonin has been shown to regulate the excitatory/inhibitory (EI) balance in visual cortical networks (Moreau et al., 2010; Carlos-Lima et al., 2023). Serotonin binds to a family of receptors that include G-protein-coupled receptors (5-HT1, 5-HT2, 5-HT4, 5-HT5, 5-HT6, and 5-HT7) (Andrade, 1998; Bockaert et al., 2006; Hannon and Hoyer, 2008; Petelák et al., 2023) and ligand-gated ion channels (5-HT3) (Kawa, 1994; Kaneez and White, 2004). The 5-HT1 family of receptors is linked to G_i/o_, therefore, activation of these receptors decreases neuronal activity (Mannoury la Cour et al., 2001; Hannon and Hoyer, 2008). The family of 5-HT2 receptors is preferentially coupled to G_q/11,_ while 5-HT4, 5-HT6, and 5-HT7 are preferentially coupled to G_s_ protein receptors. Activation of G_q/11_ and G_s_ enhances neuronal activity (Santana et al., 2004; Tang and Trussell, 2017). The serotonergic receptors previously reported in the IC are 5-HT1_A/B_, 5-HT2_A/B/C_, 5-HT3, and 5-HT7 (Hurley, 2006; Wang et al., 2008; Miko and Sanes, 2009; Ramsey et al., 2010; Papesh and Hurley, 2012). However, previous studies have used pharmacological and/or immunohistochemical approaches, which lack sufficient specificity due to the high similarity of serotonergic receptors.

Given that serotonin differently modulates distinct classes of neurons in other brain regions, such as the hippocampus (Andrade and Nicoll, 1987), the dorsal cochlear nucleus (Tang and Trussell, 2017), and the prefrontal cortex (Villalobos et al., 2005), the effect of serotonin is likely determined by the receptors expressed by classes of IC neurons. Since serotonin is a major regulator of EI balance, here we hypothesized that GABAergic and glutamatergic IC neurons differentially express serotonergic receptors. To test this hypothesis, we performed in situ hybridization targeting GABAergic and glutamatergic IC neurons, as well as six different subtypes of metabotropic serotonin receptors. Our data show that the majority of GABAergic IC neurons express excitatory serotonergic receptors (5-HT2_A_, 5-HT2_C,_ and 5-HT7), whereas glutamatergic neurons primarily express the inhibitory serotonergic receptors (5-HT1_A_ and 5-HT1_B_). Interestingly, we find little to no expression of 5-HT2_B_ in the IC. Together, our data suggest an inhibitory net effect of serotonin in the IC, poising serotonin as a major regulator of EI balance.

## Materials and Methods

### Animals

All experiments followed the National Institutes of Health’s Guide for the Care and Use of Laboratory Animals and received approval from the University of Texas at San Antonio (approval MU132). Mice had ad libitum access to food and water and were maintained on a 12h day/night cycle. Both male and female C57BL/6J mice were included in all experiments and obtained from The Jackson Laboratory (strain #000664). Since C57BL/6J mice are subject to accelerated age-related hearing loss (Noben-Trauth et al., 2003), experiments were performed in mice aged P45 – P60.

### Fluorescence in-situ hybridization assay

In situ hybridization was performed using RNAScope Multiplex Fluorescence V2 kit (Advanced Cell Diagnostics, catalog # 320850 (Wang et al., 2012). The assay was performed as described in our previous studies (Silveira et al., 2023, 2024). To collect the brains, mice were deeply anesthetized with isoflurane. Brains were quickly dissected from 10 mice (5 males and 5 females C57BL/6J aged P45 – P60), frozen in dry ice, and maintained at-80 °C until slicing. Before slicing, brains were equilibrated at-20 °C, and 15 µm sections were collected using a cryostat and mounted on Superfrost Plus slides (Fisher Scientific, catalog # 22037246). Slices were fixed in 10% neutral buffered formalin (Sigma-Aldrich, catalog # HT501128) for 1h, dehydrated in increasing concentrations of ethanol, followed by the drawing of a hydrophobic barrier around the sections. Hydrogen peroxide was used to block endogenous peroxidase for 15min at room temperature. To identify possible colocalization of serotonergic receptors (Figure 7), we used different combinations of probes for hybridization: (1) *Vgat*, *Htr1b,* and *Htr2a;* (2) *Vgat*, *Htr1a*, and *Htr1b*; (3) *Vgat*, *Htr1a*, and *Htr2c*; (4) *Vgat*, *Htr2a*, and *Htr2c;* (5) *Vgat*, *Htr7* and *Htr2b;* (6) *Vglut2*, *Htr1a*, and *Htr1b;* (7) *Vglut2*, *Htr2a*, and *Htr2c*; (8) *Vglut2*, *Htr7*, and *Htr2b*. All probes, positive controls, and negative controls were incubated for 2 hours, followed by the amplification (AMP 1 – 3) step. Signals were developed using the appropriate Horseradish Peroxidase. After that, opal dyes (1:1000) were assigned for each channel: *Vgat* and *Vglut2* expression were identified by Opal 520 (Akoya Bioscience, catalog # FP1487001KT), serotonergic receptors were identified by either Opal 570 (Akoya Bioscience, catalog # FP1488001KT) or 690 (Akoya Bioscience, catalog # FP1497001KT). Following staining with DAPI, slices were coverslipped using ProLong Gold antifade mountant (Fisher Scientific, catalog # P36934). Images were collected using a Zeiss 710 confocal microscope within two to three weeks after the assay. Representative sections (including caudal, mid-rostrocaudal, and rostral, 2-5 sections per mouse) were imaged using a 40X objective at 1-4 µm depth intervals.

### In-situ hybridization analysis

Quantification was performed either manually using FIJI (Image J, National Institutes of Health (Schindelin et al., 2012)) or semi-automatically using QuPath 0.5.1. All *Vgat^+^* and *Vglut2^+^* cells were counted in every slice. For both approaches, channels of each color were quantified separately to avoid bias. For the quantification using QuPath, we used the “cell positive detection” tool. A grid encompassing the entire IC was overlaid on each image, and 6-7 representative regions of the grid were selected to establish the optimal parameters for detecting positive cells. Next, a customized script was used to identify all positive cells in the IC. Importantly, all slices were subsequently reviewed manually to confirm accurate detection and to remove any falsely identified cells. Cells mislabeled using the automated approach were corrected in the final manual analysis. The central nucleus (ICc) and the shell IC (ICx) subdivisions were differentiated using separate tissue series stained for GAD67 and GlyT2 proteins (Choy Buentello et al., 2015; Silveira et al., 2020).

## Statistics

Statistical analyses were performed using Igor Pro 9 (Wavemetrics). For the comparison of the distribution of GABAergic and glutamatergic neurons expressing a serotonergic receptor across the whole IC, central nucleus of the IC, and shell IC, we used Welch’s t-test. The significance level (α) was adjusted to account for multiple comparisons using Bonferroni correction (α: 0.05/3 = 0.017). For the comparison between the distribution of *Vgat^+^* or *Vglut2^+^* expressing a serotonergic receptor in the central nucleus *vs* shell IC, we used Welch’s t-test, and neurons were considered differentially distributed when P < 0.05. Data are shown as means ± SD. Boxplots represent median, 25^th^ and 75^th^ percentile (box), and 9^th^ and 91^st^ percentile (whiskers).

## Results

The goal of this study was to determine the subtypes of metabotropic serotonergic receptors expressed by GABAergic and glutamatergic neurons in the IC. We focused on six metabotropic serotonergic receptors that have been previously suggested to modify how neurons encode auditory stimuli in the IC using pharmacological approaches (Hurley, 2006; Bohorquez and Hurley, 2009; Papesh and Hurley, 2012). For each serotonergic receptor, at least three mice of either sex were used for in situ hybridization.

### The 5-HT1_A_ (Htr1a) receptor is primarily expressed by IC glutamatergic neurons

The 5-HT1_A_ receptor is a G_i/o_ G-protein receptor that inhibits neuronal activity by activating G protein-gated inwardly rectifying potassium channels (GIRK) (Bockaert et al., 2006). Activation of the 5-HT1_A_ receptor suppresses responses to broadband vocalizations and alters responses to tone and frequency-modulated (FM) sweeps in the IC (Ramsey et al., 2010; Gentile Polese et al., 2021). However, the distribution of this receptor in the IC and which neurons express this receptor are unknown.

In the first essay, we used probes to identify *Vgat* (GABAergic marker) and *Htr1a.* For that, we used thirteen brain slices from three males (P45) and one female (P59) C57BL/6 mice. Our data show that 4.6% (298 out of 6967) of *Vgat^+^* neurons express *Htr1a* (**Figure 1A-D, I, Table 1**). In a separate assay, we used probes to identify *Vglut2* (glutamatergic marker) and *Htr1a.* We found ∼27% (20782 out of 78046) of IC *Vglut2^+^* neurons express *Htr1a* (**Figure 1E-H, J, Table 2**). The percentage of glutamatergic neurons expressing *Htr1a* was higher than that of GABAergic neurons across the whole IC, as well as within the central nucleus and shell subdivisions of the IC. (**Figure 1K**, IC: 4.6 ± 1.4% of *Vgat^+^Htr1a^+^ vs* 26.6 ± 7.0% of *Vglut^+^Htr1a^+^*, Welch’s t test, α = 1 x 10^-9^; ICc: 3.7 ± 2.3% of *Vgat^+^Htr1a^+^ vs* 26.2 ± 7.0% of *Vglut2^+^Htr1a^+^*, Welch’s t test, α = 2 x 10^-8^; ICx: 5.6 ± 1.6% of *Vgat^+^Htr1a^+^ vs* 27.2 ± 6.1% of *Vglut2^+^Htr1a^+^*, Welch’s t test, α = 1 x 10^-10^).

**Figure 1.**
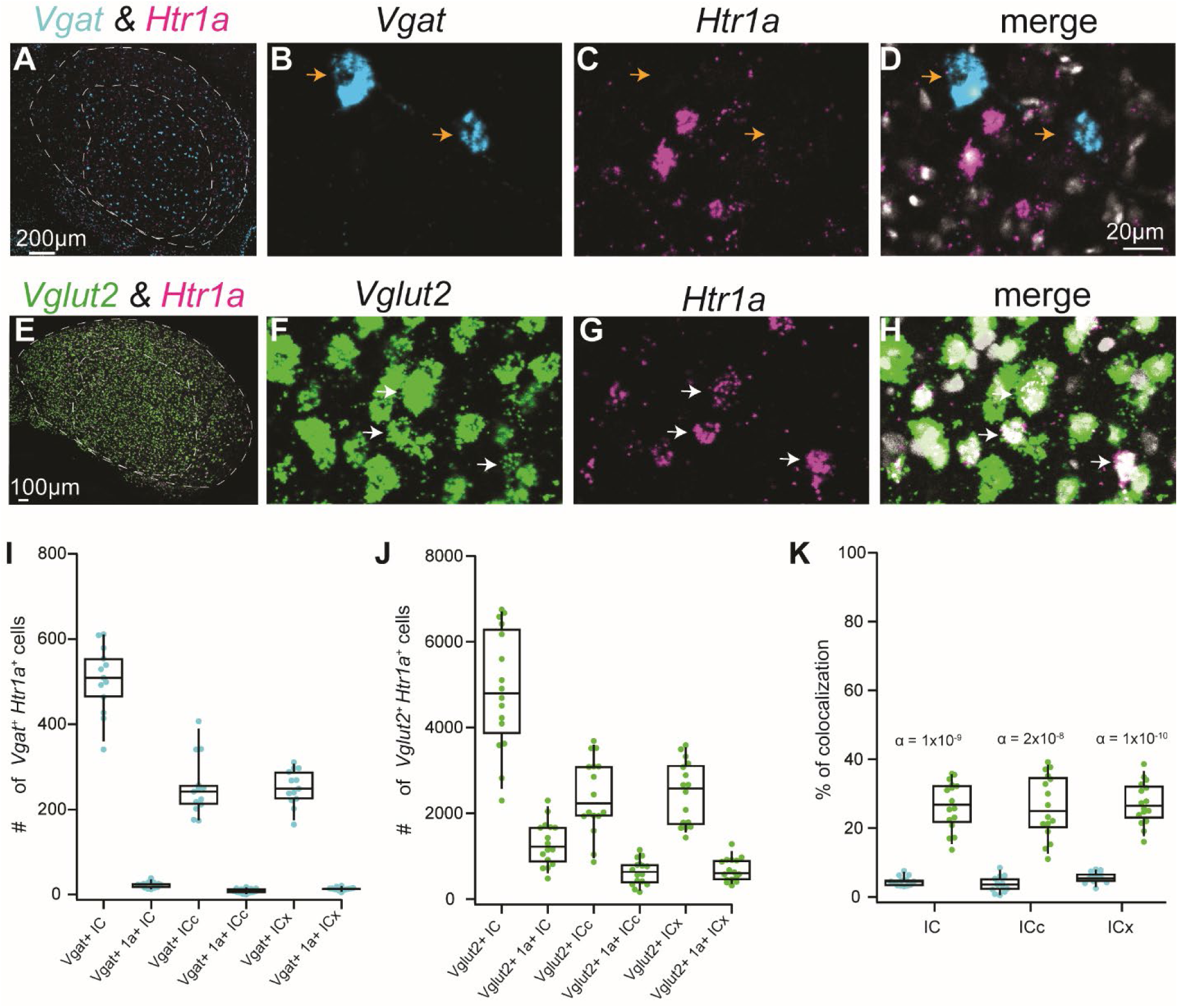
The 5-HT1A (Htr1a) receptor is primarily expressed by IC glutamatergic neurons. Fluorescent in situ hybridization was used to determine the expression patterns of Vgat, Vglut2, and Htr1a in the C57BL/6J mice. **A.** Coronal IC section showing the distribution of Vgat (cyan) and Htr1a (magenta) in the IC. **B-D.** High magnification confocal images showing that Vgat+ neurons rarely colabel with Htr1a, as shown by orange arrows. DAPI is shown in white. **E.** Coronal IC section showing the distribution of Vglut2 (green) and Htr1a (magenta) in the IC. **F-H.** High magnification confocal images showing that Vglut2^+^ neurons often colabel with Htr1a, as shown by white arrows. **I.** Boxplot showing the number of Vgat^+^ neurons and Vgat^+^ neurons that express the Htr1a in the IC, as well as in the central nucleus of the IC and shell IC. **J.** Boxplot showing the number of Vglut2^+^ neurons and Vglut2+ neurons that express the Htr1a in the IC, as well as in the central nucleus of the IC and shell IC. **K.** Comparison of the percentage of Vgat^+^ neurons expressing Htr1a (cyan dots) vs the percentage of Vglut2^+^ neurons expressing Htr1a (green dots) in the whole IC, central nucleus of the IC, and shell IC. α-values from Welch’s t-tests are atop each plot, showing the difference between the percentage of Vgat^+^ vs Vglut2^+^expressing Htr1a.

**Table 1.**
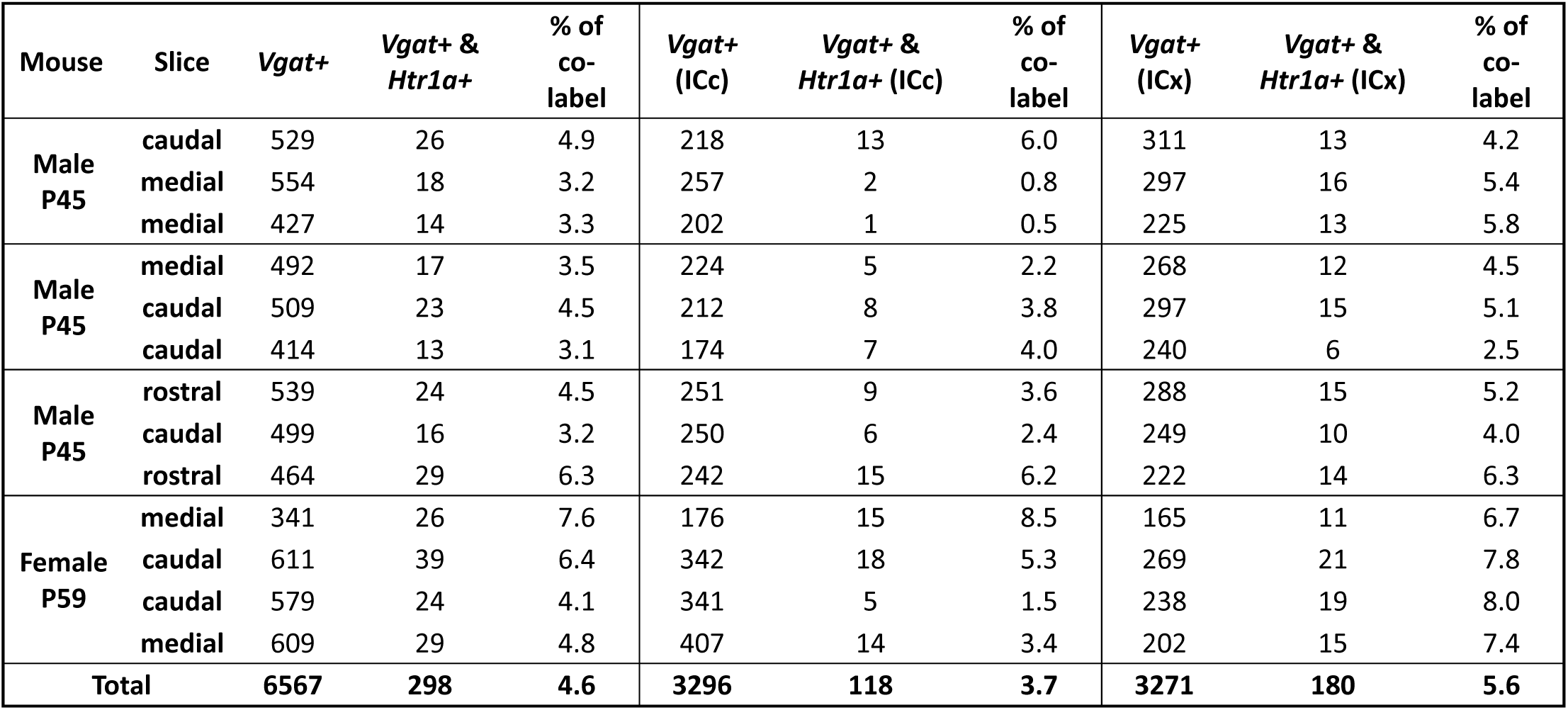
Nearly 4.5% of *Vgat^+^* neurons express *Htr1a*.

**Table 2.** Nearly 27% of *Vglut2^+^* neurons express *Htr1a*.

| Mouse | Slice | *Vglut2*<sup>+</sup> | *Vglut2*<sup>+</sup> & *Htr1a*<sup>+</sup> | % of co-label | *Vglut2*<sup>+</sup> (ICc) | *Vglut2*<sup>+</sup> & *Htr1a*<sup>+</sup> (ICc) | % of co-label | *Vglut2*<sup>+</sup> (ICx) | *Vglut2*<sup>+</sup> & *Htr1a*<sup>+</sup> (ICx) | % of co-label |
| --- | --- | --- | --- | --- | --- | --- | --- | --- | --- | --- |
| Male P45 | caudal | 2820 | 717 | 25.4 | 1038 | 221 | 21.3 | 1782 | 496 | 27.8 |
|  | medial | 4907 | 1678 | 34.2 | 2020 | 762 | 37.7 | 2887 | 916 | 31.7 |
|  | rostral | 4689 | 1665 | 35.5 | 2023 | 794 | 39.2 | 2666 | 871 | 32.7 |
| Male P45 | caudal | 3590 | 1156 | 32.2 | 1931 | 579 | 30.0 | 1659 | 577 | 34.8 |
|  | caudal | 4090 | 1050 | 25.7 | 1592 | 425 | 26.7 | 2498 | 625 | 25.0 |
|  | caudal | 2298 | 479 | 20.8 | 864 | 164 | 19.0 | 1434 | 315 | 22.0 |
| Male P45 | caudal | 5106 | 1659 | 32.5 | 1947 | 685 | 35.2 | 3159 | 974 | 30.8 |
|  | medial | 6176 | 1722 | 27.9 | 3514 | 816 | 23.2 | 2662 | 906 | 34.0 |
|  | rostral | 6582 | 2042 | 31.0 | 3090 | 1145 | 37.1 | 3492 | 897 | 25.7 |
| Female P59 | caudal | 3625 | 812 | 22.4 | 1939 | 411 | 21.2 | 1686 | 401 | 23.8 |
|  | medial | 6669 | 913 | 13.7 | 3079 | 340 | 11.0 | 3590 | 573 | 16.0 |
|  | medial | 6748 | 1237 | 18.3 | 3685 | 564 | 15.3 | 3063 | 585 | 19.1 |
| Female P60 | caudal | 4509 | 1428 | 31.7 | 2851 | 975 | 34.2 | 1658 | 453 | 27.3 |
|  | medial | 6415 | 2296 | 35.8 | 3089 | 1012 | 32.8 | 3326 | 1284 | 38.6 |
|  | medial | 4224 | 725 | 17.2 | 2436 | 340 | 14.0 | 1788 | 385 | 21.5 |
|  | rostral | 5598 | 1291 | 23.1 | 3519 | 777 | 22.1 | 2079 | 514 | 24.7 |
| Total |  | 78046 | 20782 | 26.7 | 38617 | 10010 | 26.2 | 39429 | 10772 | 27.2 |

When analyzing the distribution of GABAergic neurons expressing *Htr1a* across the central nucleus and the shell, we found that even though a small percentage of GABAergic neurons express *Htr1a, Vgat^+^Htr1a^+^* neurons were more likely to be present in the shell (ICc: 3.7 ± 2.3% *vs* ICx: 5.6 ± 1.6% of *Vgat^+^Htr1a^+^.* Welch’s *t test,* P = 0.024). On the other hand, there was no statistical difference in the percentage of glutamatergic neurons expressing *Htr1a* in the central IC and the shell IC (ICc: 26.2 ± 9.1% *vs* ICx: 27.2 ± 6.1% of *Vgat^+^Htr1a^+^.* Welch’s *t test,* P = 0.727).

### Many IC glutamatergic neurons express the 5-HT1_B_ (Htr1b) receptor

Activation of the 5-HT1_B_ receptor inhibits neuronal activity in various brain regions, including the substantia nigra and globus pallidus, where 5-HT1_B_ modulates GABAergic transmission (Johnson et al., 1992; Pommer et al., 2021), as well as in the bed nucleus of the stria terminalis, in which 5-HT1_B_ inhibits glutamatergic transmission (Guo and Rainnie, 2010). In the IC, it has been suggested that activation of the 5-HT1_B_ receptor alters the frequency tuning of neurons by modulating GABA release (Hurley et al., 2008). Our in-situ hybridization data show that only 3.5% (318 out of 9000) of GABAergic neurons express the *Htr1b* receptor (**Figure 2A-D, I, Table 3**). On the other hand, ∼27% (21244 out of 78046) of glutamatergic neurons express the *Htr1b* receptor (**Figure 2E-H, J, Table 4**). The percentage of glutamatergic neurons expressing *Htr1b* was higher compared to GABAergic neurons. (**Figure 2K**, IC: 3.5 ± 1.5% of *Vgat^+^Htr1b^+^ vs* 26.8 ± 6.6% of *Vglut2^+^Htr2b^+^*, Welch’s *t test,* α = 1 x 10^-10^; ICc: 2.1 ± 1.2% of *Vgat^+^Htr1b^+^ vs* 24.1 ± 7.7% of *Vglu2t^+^Htr1b^+^*, Welch’s *t test,* α = 5 x 10^-9^; ICx: 5.5 ± 2.7% of *Vgat^+^Htr1b^+^ vs* 29.9 ± 7.0% of *Vglu2t^+^Htr1b^+^*, Welch’s *t test,* α = 5 x 10^-11^).

**Figure 2.**
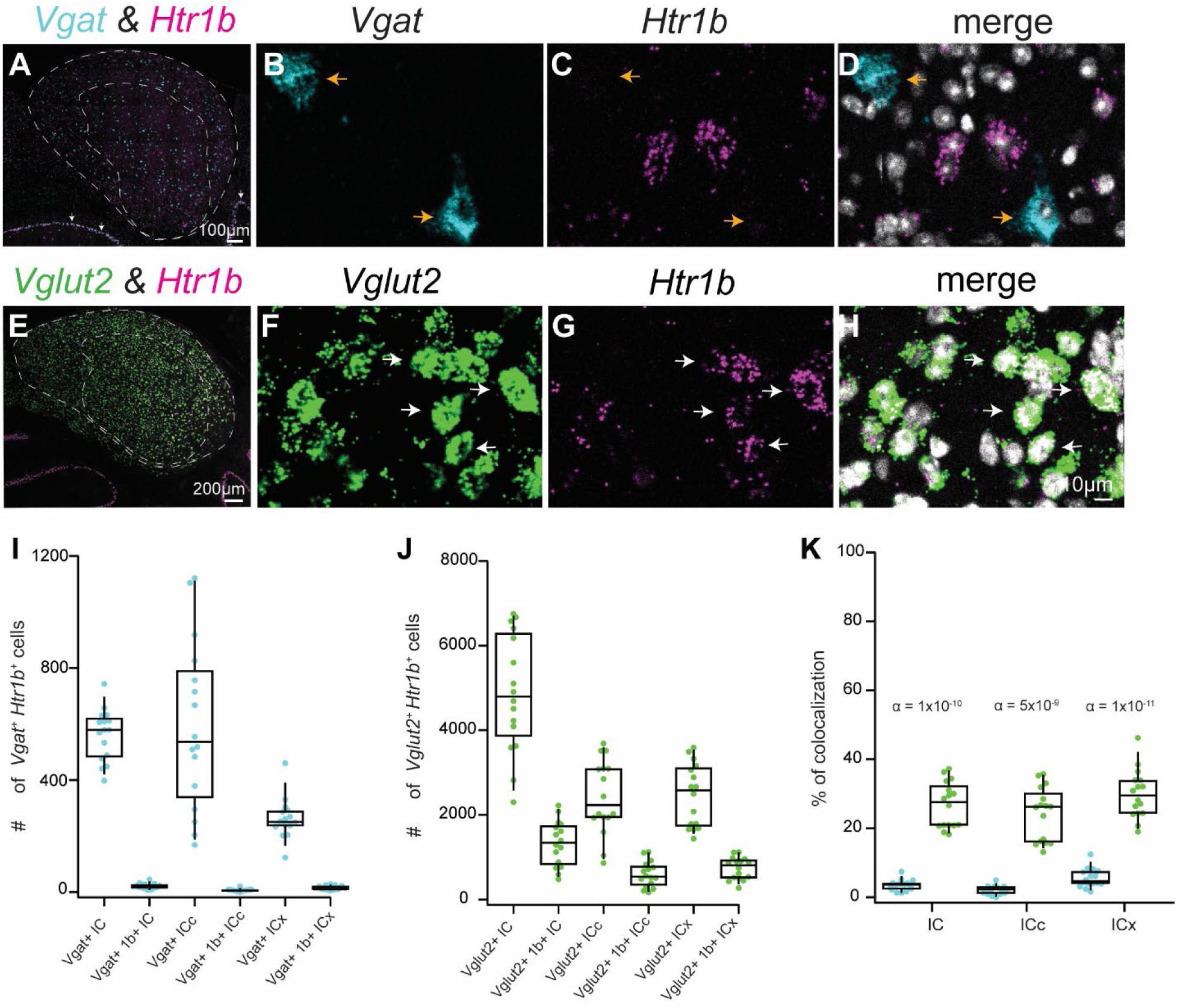
Many IC glutamatergic neurons express the 5-HT1B (Htr1b) receptor. Fluorescent in situ hybridization was used to determine the expression patterns of Vgat, Vglut2, and Htr1b in the C57BL/6J mice. **A.** Coronal IC section showing the distribution of Vgat (cyan) and Htr1b (magenta) in the IC. White arrows show the colocalization between Purkinje cells in the cerebellum and Htr1b. **B-D.** High magnification confocal images showing that Vgat+ neurons rarely colabel with Htr1b, as shown by orange arrows. DAPI is shown in white. **E.** Coronal IC section showing the distribution of Vglut2 (green) and Htr1b (magenta) in the IC. **F-H.** High magnification confocal images showing that Vglut2+ neurons often colabel with Htr1b, as shown by white arrows. **I.** Boxplot showing the number of Vgat^+^ neurons and Vgat+ neurons that express the Htr1b in the IC, as well as in the central nucleus of the IC and shell IC. **J.** Boxplot showing the number of Vglut2+ neurons and Vglut2+ neurons that express the Htr1b in the IC, as well as in the central nucleus of the IC and shell IC. **K.** Comparison of the percentage of Vgat+ neurons expressing Htr1b (cyan dots) vs the percentage of Vglut2^+^ neurons expressing Htr1b (green dots) in the whole IC, central nucleus of the IC, and shell IC. α-values from Welch’s t-tests are atop each plot, showing the difference between the percentage of Vgat^+^ vs Vglut2^+^expressing Htr1b.

**Table 3.** Nearly 3.5% of *Vgat^+^* neurons express *Htr1b*.

| Mouse | Slice | *Vgat*<sup>+</sup> | *Vgat*<sup>+</sup> & *Htr1b*<sup>+</sup> | % of co-label | *Vgat*<sup>+</sup> (ICc) | *Vgat*<sup>+</sup> & *Htr1b*<sup>+</sup> (ICc) | % of co-label | *Vgat*<sup>+</sup> (ICx) | *Vgat*<sup>+</sup> & *Htr1b*<sup>+</sup> (ICx) | % of co-label |
| --- | --- | --- | --- | --- | --- | --- | --- | --- | --- | --- |
| Male P45 | caudal | 631 | 8 | 1.3 | 345 | 1 | 0.3 | 286 | 7 | 2.4 |
|  | caudal | 488 | 18 | 3.7 | 249 | 6 | 2.4 | 239 | 12 | 5.0 |
|  | medial | 657 | 23 | 3.5 | 396 | 7 | 1.8 | 261 | 16 | 6.1 |
|  | rostral | 606 | 8 | 1.3 | 302 | 3 | 1.0 | 304 | 5 | 1.6 |
| Male P45 | caudal | 398 | 14 | 3.5 | 194 | 6 | 3.1 | 204 | 8 | 3.9 |
|  | caudal | 441 | 16 | 3.6 | 192 | 6 | 3.1 | 249 | 10 | 4.0 |
|  | medial | 569 | 14 | 2.5 | 317 | 4 | 1.3 | 252 | 10 | 4.0 |
|  | rostral | 448 | 18 | 4.0 | 325 | 9 | 2.8 | 123 | 9 | 7.3 |
| Male P45 | caudal | 579 | 13 | 2.2 | 334 | 3 | 0.9 | 245 | 10 | 4.1 |
|  | caudal | 476 | 18 | 3.8 | 242 | 8 | 3.3 | 234 | 10 | 4.3 |
|  | medial | 632 | 32 | 5.1 | 302 | 7 | 2.3 | 330 | 25 | 7.6 |
|  | rostral | 533 | 9 | 1.7 | 241 | 0 | 0.0 | 292 | 9 | 3.1 |
| Female P59 | caudal | 611 | 30 | 4.9 | 342 | 8 | 2.3 | 269 | 22 | 8.2 |
|  | caudal | 579 | 24 | 4.1 | 341 | 4 | 1.2 | 238 | 20 | 8.4 |
|  | medial | 743 | 28 | 3.8 | 283 | 9 | 3.2 | 460 | 28 | 6.1 |
|  | medial | 609 | 45 | 7.4 | 407 | 20 | 4.9 | 202 | 25 | 12.4 |
| Total |  | 9000 | 318 | 3.5 | 4812 | 101 | 2.1 | 4188 | 226 | 5.5 |

**Table 4.** Nearly 27% of *Vglut2^+^* neurons express *Htr1b*.

| Mouse | Slice | *Vglut2*<sup>+</sup> | *Vglut2*<sup>+</sup> & *Htr1b*<sup>+</sup> | % of co-label | *Vglut2*<sup>+</sup> (ICc) | *Vglut2*<sup>+</sup> & *Htr1b*<sup>+</sup> (ICc) | % of co-label | *Vglut2*<sup>+</sup> (ICx) | *Vglut2*<sup>+</sup> & *Htr1b*<sup>+</sup> (ICx) | % of co-label |
| --- | --- | --- | --- | --- | --- | --- | --- | --- | --- | --- |
| Male P45 | caudal | 2820 | 591 | 21.0 | 1038 | 168 | 16.2 | 1782 | 429 | 24.1 |
|  | medial | 4907 | 1779 | 36.3 | 2020 | 667 | 33.0 | 2887 | 1112 | 38.5 |
|  | rostral | 4689 | 1614 | 34.4 | 2023 | 715 | 35.3 | 2666 | 903 | 33.9 |
| Male P45 | caudal | 3590 | 745 | 20.8 | 1931 | 295 | 15.3 | 1659 | 450 | 27.1 |
|  | caudal | 4090 | 767 | 18.8 | 1592 | 251 | 15.8 | 2498 | 516 | 20.7 |
|  | caudal | 2298 | 476 | 20.7 | 864 | 204 | 23.6 | 1434 | 272 | 19.0 |
| Male P45 | caudal | 5106 | 1531 | 30.0 | 1947 | 554 | 28.5 | 3159 | 977 | 30.9 |
|  | medial | 6176 | 1820 | 29.5 | 3514 | 918 | 26.1 | 2662 | 902 | 33.9 |
|  | rostral | 6582 | 2221 | 33.7 | 3090 | 1104 | 35.7 | 3492 | 1117 | 32.0 |
| Female P59 | caudal | 3625 | 1122 | 31.0 | 1939 | 509 | 26.3 | 1686 | 613 | 36.4 |
|  | medial | 6669 | 1393 | 20.9 | 3079 | 518 | 16.8 | 3590 | 875 | 24.4 |
|  | medial | 6748 | 1228 | 18.2 | 3685 | 483 | 13.1 | 3063 | 745 | 24.3 |
| Female P60 | caudal | 4509 | 1289 | 28.6 | 2851 | 756 | 26.5 | 1658 | 533 | 32.1 |
|  | medial | 6415 | 1705 | 26.6 | 3089 | 826 | 26.7 | 3326 | 879 | 26.4 |
|  | medial | 4224 | 882 | 20.9 | 2436 | 379 | 15.6 | 1788 | 503 | 28.1 |
|  | rostral | 5598 | 2081 | 37.2 | 3519 | 1121 | 31.9 | 2079 | 960 | 46.2 |
| Total |  | 78046 | 21244 | 26.8 | 38617 | 9468 | 24.1 | 39429 | 11786 | 29.9 |

Even though there was a small percentage of GABAergic neurons expressing *Htr1b,* these neurons were more likely to be present in the shell IC (2.1 ± 1.2% of *Vgat^+^Htr1b^+^* neurons in the ICc *vs* 5.5 ± 2.7% of *Vgat^+^Htr1b^+^* neurons in the ICx. Welch’s *t test,* P = 0.0001). Similarly, glutamatergic neurons expressing *Htr1b* were more concentrated in the shell IC (24.1 ± 7.7% of *Vglut2^+^Htr1b^+^* neurons in the ICc *vs* 29.9 ± 7.0% of *Vglut2^+^Htr1b^+^* neurons in the ICx. Welch’s *t test,* P = 0.036).

### Over half of IC GABAergic neurons express the 5-HT2_A_ (Htr2a) receptor

The family of 5-HT2 receptors is preferentially coupled to G_q/11_. Thereby, activation of these receptors enhances neuronal activity (Bockaert et al., 2006). The 5-HT2_A_ serotonergic receptor has been suggested to modulate GABAergic transmission in the IC (Wang et al., 2008). However, since the IC integrates many GABAergic inputs, it is unknown whether this receptor is expressed by GABAergic neurons outside the IC and/or by local GABAergic IC neurons. We found that over half of IC *Vgat^+^* neurons express the *Htr2a* receptor (52.2%, 5319 out of 10124 neurons, **Figure 3A-D, I, Table 5**). Expression of *Htr2a* was rarely seen in glutamatergic IC neurons since only 2.7% of glutamatergic neurons express this receptor (**Figure 3E-H, J, Table 6**). The percentage of GABAergic neurons expressing *Htr2a^+^* was higher than the percentage of glutamatergic neurons expressing *Htr2a^+^* across the whole IC, central IC, and shell IC (**Figure 3K**, IC: 52.2 ± 8.5% of *Vgat^+^Htr2a^+^ vs* 2.7 ± 0.8% of *Vglut2^+^Htr2a^+^*, Welch’s *t test,* α = 1 x 10^-15^; ICc: 61.1 ± 14.3% of *Vgat^+^Htr2a^+^ vs* 3.0 ± 1.3% of *Vglu2t^+^Htr2a^+^*, Welch’s *t test,* α = 6 x 10^-13^; ICx: 41.3 ± 8.5% of *Vgat^+^Htr2a^+^ vs* 2.4 ± 0.7% of *Vglu2t^+^Htr2a^+^*, Welch’s *t test,* α = 7 x 10^-14^).

**Figure 3.**
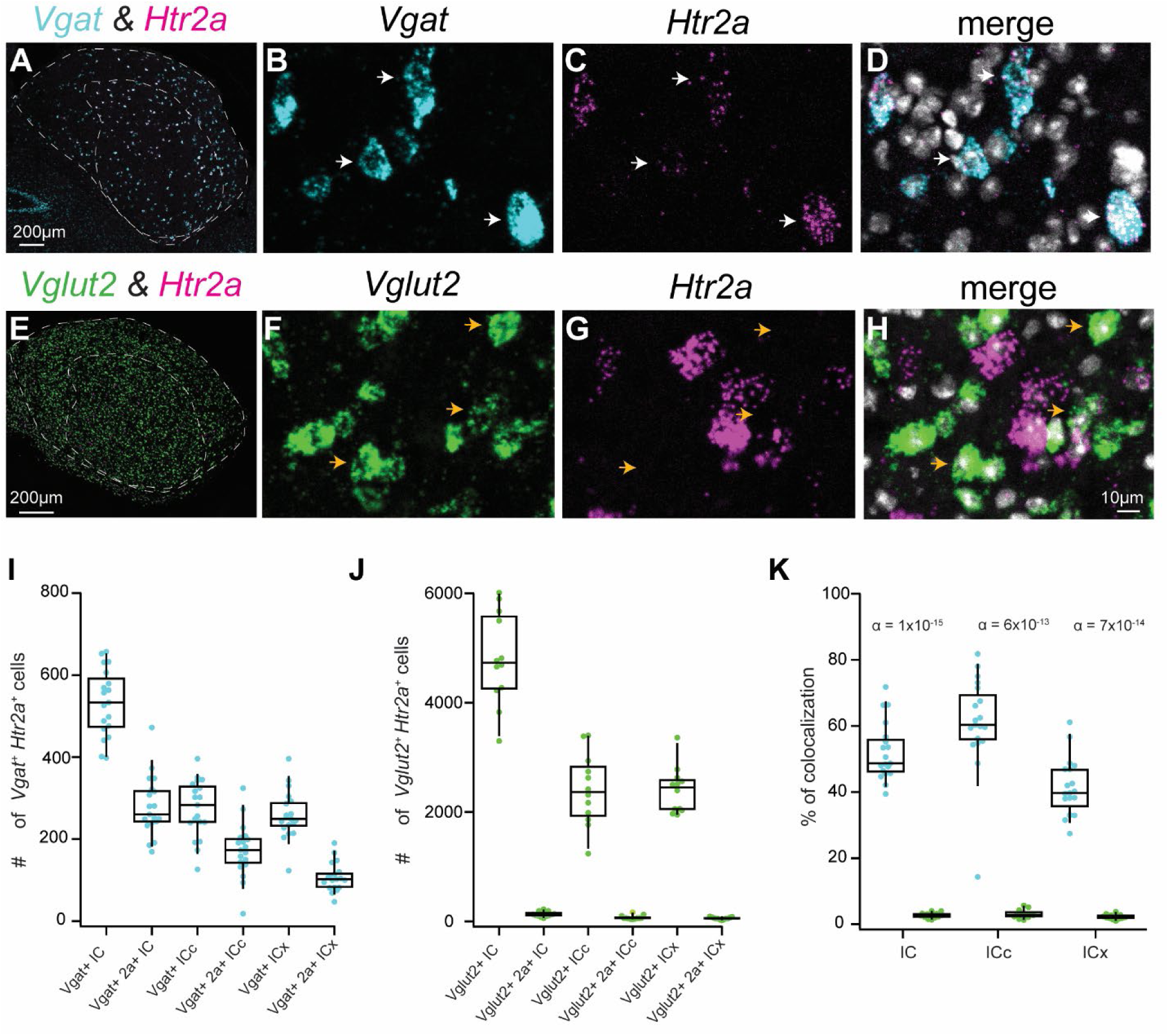
Over half of IC GABAergic neurons express the 5-HT2A (Htr2a) receptor. Fluorescent in situ hybridization was used to determine the expression patterns of Vgat, Vglut2, and Htr2a in the C57BL/6J mice. **A.** Coronal IC section showing the distribution of Vgat (cyan) and Htr2a (magenta) in the IC. **B-D.** High magnification confocal images showing that Vgat^+^ neurons highly colabel with Htr2a, as shown by white arrows. **E.** Coronal IC section showing the distribution of Vglut2 (green) and Htr2a (magenta) in the IC. **F-H.** High magnification confocal images showing that Vglut2^+^ neurons rarely colabel with Htr2a, as shown by orange arrows. **I.** Boxplot showing the number of Vgat^+^ neurons and Vgat^+^ neurons that express the Htr2a in the IC, as well as in the central nucleus of the IC and shell IC. **J.** Boxplot showing the number of Vglut2^+^ neurons and Vglut2^+^ neurons that express the Htr2a in the IC, as well as in the central nucleus of the IC and shell IC. **K.** Comparison of the percentage of Vgat^+^ \neurons expressing Htr2a (cyan dots) vs the percentage of Vglut2^+^ neurons expressing Htr2a (green dots) in the whole IC, central nucleus of the IC, and shell IC. α-values from Welch’s t-tests are atop each plot, showing the difference between the percentage of Vgat^+^ vs Vglut2^+^expressing Htr2a.

**Table 5.**
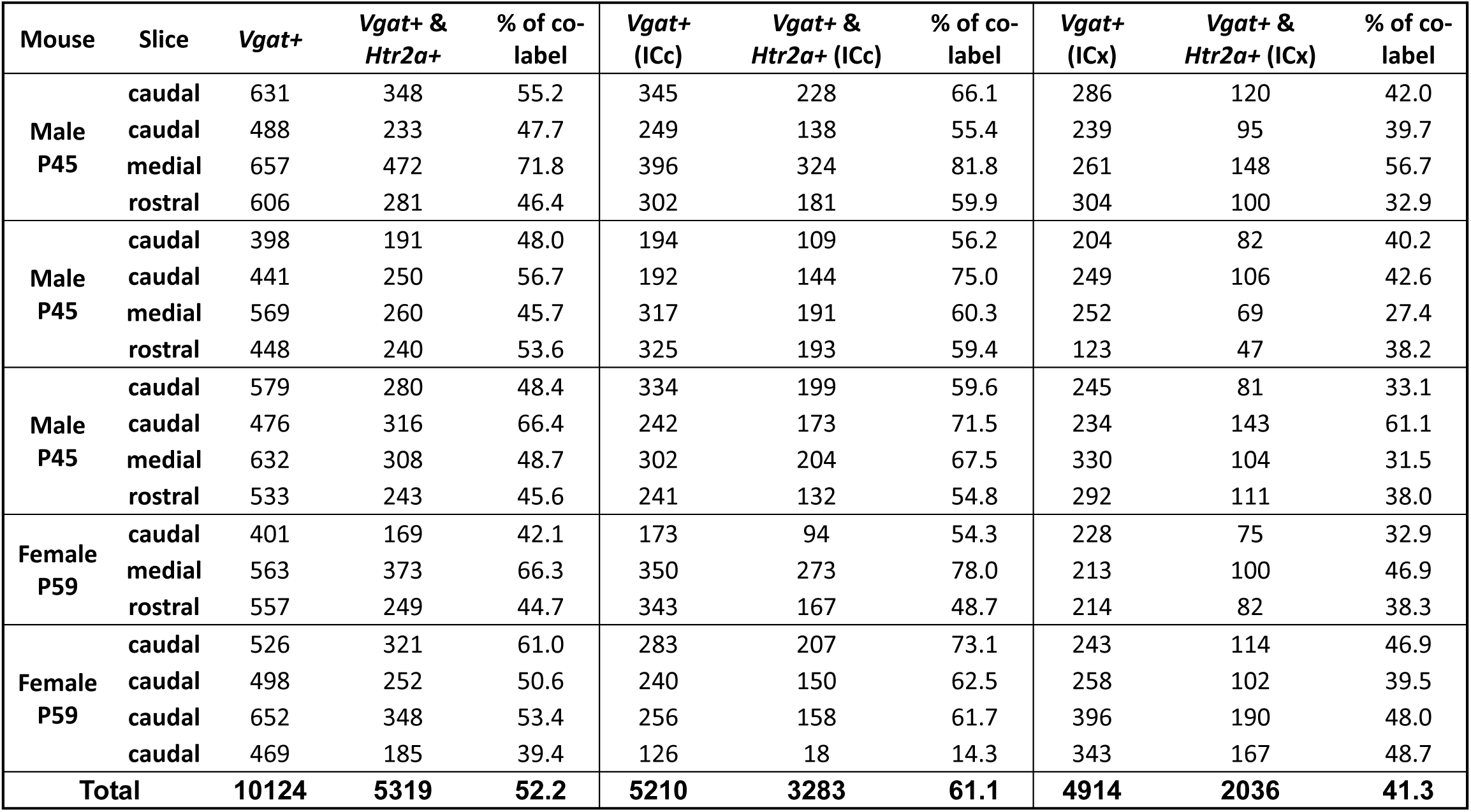
Over 50% of *Vgat^+^* neurons express *Htr2a*.

**Table 6.** Less than 3% of *Vglut2+* neurons express *Htr2a*.

| Mouse | Slice | *Vglut2*<sup>+</sup> | *Vglut2*<sup>+</sup> & *Htr2a*<sup>+</sup> | % of co-label | *Vglut2*<sup>+</sup> (ICc) | *Vglut2*<sup>+</sup> & *Htr2a*<sup>+</sup> (ICc) | % of co-label | *Vglut2*<sup>+</sup> (ICx) | *Vglut2*<sup>+</sup> & *Htr2a*<sup>+</sup> (ICx) | % of co-label |
| --- | --- | --- | --- | --- | --- | --- | --- | --- | --- | --- |
| **Male P45** | caudal | 6010 | 192 | 3.2 | 3384 | 101 | 3.0 | 2626 | 91 | 3.5 |
|  | caudal | 5900 | 81 | 1.4 | 3399 | 44 | 1.3 | 2501 | 37 | 1.5 |
|  | medial | 4690 | 129 | 2.8 | 2740 | 42 | 1.5 | 1950 | 87 | 4.5 |
| **Female P59** | caudal | 3299 | 102 | 3.1 | 1240 | 53 | 4.3 | 2059 | 49 | 2.4 |
|  | caudal | 4224 | 138 | 3.3 | 2168 | 60 | 2.8 | 2056 | 78 | 3.8 |
|  | medial | 5498 | 221 | 4.0 | 2938 | 167 | 5.7 | 2560 | 54 | 2.1 |
| **Female P59** | caudal | 4273 | 110 | 2.6 | 1767 | 59 | 3.3 | 2506 | 51 | 2.0 |
|  | caudal | 4767 | 111 | 2.3 | 1987 | 60 | 3.0 | 2780 | 51 | 1.8 |
|  | medial | 5674 | 125 | 2.2 | 2314 | 63 | 2.7 | 3360 | 62 | 1.8 |
| **Female P57** | caudal | 3827 | 59 | 1.5 | 1859 | 40 | 2.2 | 1968 | 19 | 1.0 |
|  | caudal | 4656 | 94 | 2.0 | 2625 | 61 | 2.3 | 2031 | 33 | 1.6 |
|  | medial | 4814 | 192 | 4.0 | 2418 | 125 | 5.2 | 2396 | 67 | 2.8 |
| **Total** |  | **57632** | **1554** | **2.7** | **28839** | **875** | **3.0** | **28793** | **679** | **2.4** |

GABAergic neurons expressing *Htr2a* were widely distributed throughout the IC. However, there was a higher percentage of GABAergic neurons expressing the *Htr2a* in the central IC when compared to the shell IC (61.1 ± 14.3% of *Vgat^+^Htr2a^+^* neurons in the ICc *vs* 41.2 ± 8.5% of *Vgat^+^Htr2a^+^* neurons in the ICx. Welch’s *t test,* P = 0.00001). No difference was observed in the distribution of *Vglut2^+^Htr2a^+^* neurons across IC subdivisions (ICc: 3.1 ± 1.3% *vs* ICx: 2.2 ± 0.7% of *Vglut2^+^Htr2a^+^.* Welch’s *t test,* P = 0.066).

### The 5-HT2_B_ (Htr2b) receptor is rarely expressed in the IC of young mice

The 5-HT2_B_ receptor has low expression in the brain (Bockaert et al., 2006). Expression of this receptor has been reported in the cerebellum, medial amygdala, and dorsal hypothalamus (Wainscott et al., 1993; Bockaert et al., 2006). On the other hand, this receptor is highly expressed in cardiac tissue and lungs (Launay et al., 2002; Nebigil et al., 2003; Hutcheson et al., 2011). A previous study reported that this receptor is also expressed in the IC. Interestingly, the expression was low in young mice but increased in older mice with hearing loss (Tadros et al., 2007). Here, we observed little to no expression of *Htr2b* in the IC of young mice (**Figure 4** and **Tables 7 and 8**). Future studies will be necessary to determine whether these receptors are expressed by either GABAergic and/or glutamatergic neurons in older mice.

**Figure 4.**
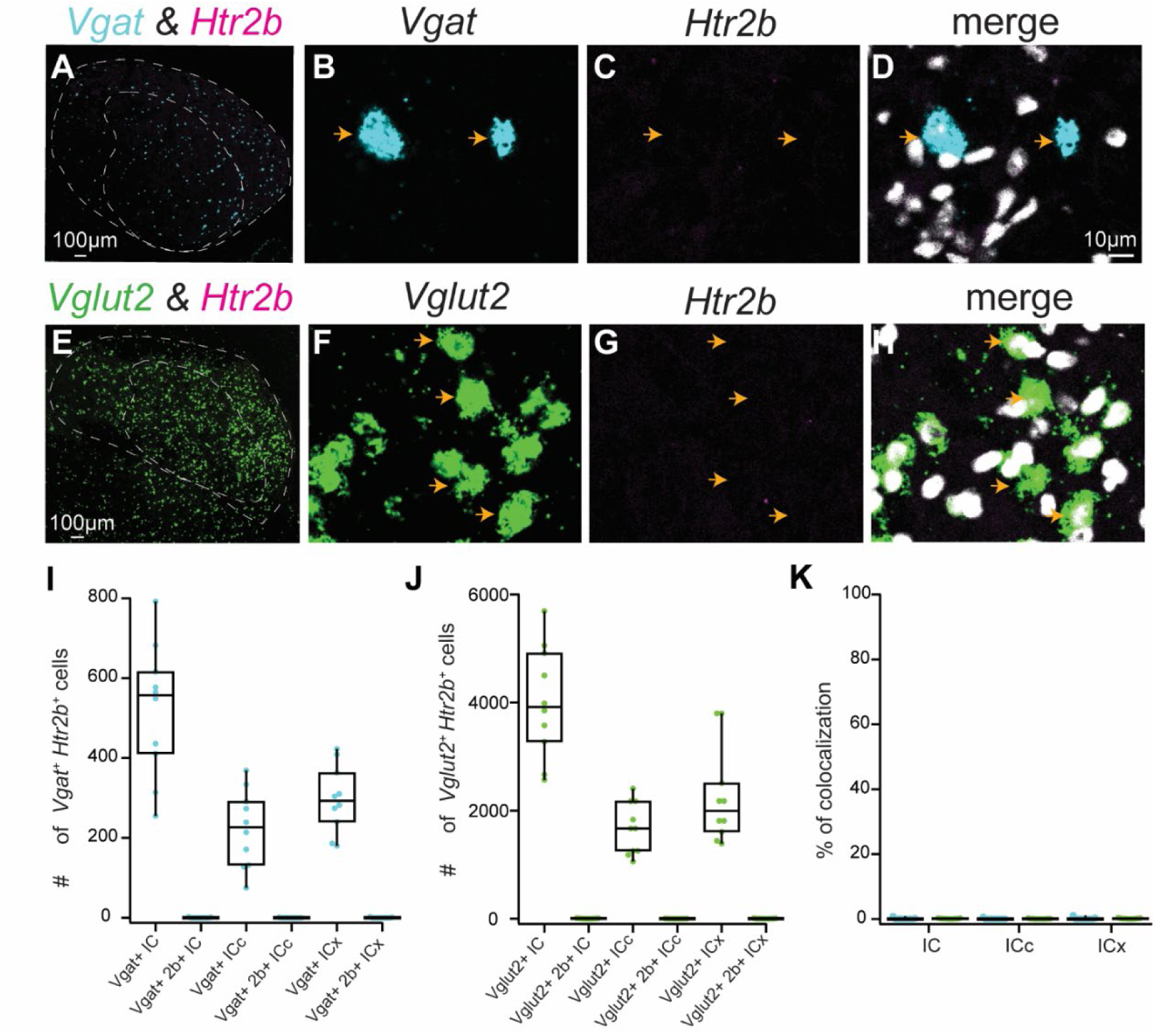
The 5-HT2B (Htr2b) receptor is rarely expressed in the IC of young mice. Fluorescent in situ hybridization was used to determine the expression patterns of Vgat, Vglut2, and Htr2b in the C57BL/6J mice. **A.** Coronal IC section showing the distribution of Vgat (cyan) and Htr2b (magenta) in the IC. **B-D.** High magnification confocal images showing that Htr2b receptors are not expressed in the IC. **E.** Coronal IC section showing the distribution of Vglut2 (green) and Htr2b (magenta) in the IC. **F-H.** High magnification confocal images showing the lack of expression of Htr2b in the IC, as shown by orange arrows. **I.** Boxplot showing the number of Vgat^+^ neurons and Vgat^+^ neurons that express the Htr2b in the IC, as well as in the central nucleus of the IC and shell IC. **J.** Boxplot showing the number of Vglut2^+^ neurons and Vglut2^+^ neurons that express the Htr2b in the IC, as well as in the central nucleus of the IC and shell IC. **K.** Boxplot showing the percentage of Vgat^+^ neurons expressing Htr2b (cyan dots) and the percentage of Vglut2^+^ neurons expressing Htr2b (green dots) in the whole IC, central nucleus of the IC, and shell IC. Given the very low number of colocalization (0.1%), statistical tests were not performed for this receptor.

**Table 7.** Htr2b is rarely expressed by GABAergic neurons.

| Mouse | Slice | Vgat+ | Vgat+ & Htr2b+ | % of co-label | Vgat+ (ICc) | Vgat+ & Htr2b+ (ICc) | % of co-label | Vgat+ (ICx) | Vgat+ & Htr2b+ (ICx) | % of co-label |
| --- | --- | --- | --- | --- | --- | --- | --- | --- | --- | --- |
| **Male P50** | caudal | 682 | 0 | 0.0 | 273 | 0 | 0.0 | 409 | 0 | 0.0 |
|  | medial | 436 | 0 | 0.0 | 132 | 0 | 0.0 | 304 | 0 | 0.0 |
|  | medial | 792 | 0 | 0.0 | 369 | 0 | 0.0 | 423 | 0 | 0.0 |
|  | rostral | 616 | 0 | 0.0 | 334 | 0 | 0.0 | 282 | 0 | 0.0 |
| **Female 59** | caudal | 549 | 0 | 0.0 | 239 | 0 | 0.0 | 310 | 1 | 0.3 |
|  | caudal | 411 | 1 | 0.2 | 171 | 1 | 0.6 | 240 | 0 | 0.0 |
|  | caudal | 255 | 2 | 0.8 | 75 | 0 | 0.0 | 180 | 2 | 1.1 |
| **Female 59** | rostral | 565 | 0 | 0.0 | 291 | 0 | 0.0 | 274 | 0 | 0.0 |
|  | rostral | 577 | 0 | 0.0 | 214 | 0 | 0.0 | 363 | 0 | 0.0 |
|  | rostral | 314 | 0 | 0.0 | 128 | 0 | 0.0 | 186 | 0 | 0.0 |
| **Total** |  | **5197** | **3** | **0.1** | **2226** | **1** | **0.1** | **2971** | **3** | **0.1** |

**Table 8.** Htr2b is rarely expressed by glutamatergic neurons.

| Mouse | Slice | Vglut2+ | Vglut2+ & Htr2b+ | % of co-label | Vglut2+ (ICc) | Vglut2+ & Htr2b+ (ICc) | % of co-label | Vglut2+ (ICx) | Vglut2+ & Htr2b+ (ICx) | % of co-label |
| --- | --- | --- | --- | --- | --- | --- | --- | --- | --- | --- |
| **Male P50** | caudal | 2668 | 0 | 0.0 | 1059 | 0 | 0.0 | 1609 | 0 | 0.0 |
|  | medial | 4915 | 6 | 0.1 | 2407 | 3 | 0.1 | 2508 | 3 | 0.1 |
|  | medial | 3276 | 4 | 0.1 | 1834 | 2 | 0.1 | 1442 | 2 | 0.1 |
|  | rostral | 2571 | 3 | 0.1 | 1182 | 0 | 0.0 | 1389 | 3 | 0.2 |
| **Male P50** | medial | 3987 | 10 | 0.3 | 2175 | 7 | 0.3 | 1812 | 3 | 0.2 |
|  | medial | 3851 | 10 | 0.3 | 1672 | 2 | 0.1 | 2179 | 8 | 0.4 |
|  | rostral | 5055 | 3 | 0.1 | 1256 | 1 | 0.1 | 3799 | 2 | 0.1 |
| **Female P45** | caudal | 3577 | 0 | 0.0 | 2175 | 0 | 0.0 | 1812 | 0 | 0.0 |
|  | caudal | 5694 | 12 | 0.2 | 1256 | 0 | 0.0 | 3799 | 12 | 0.3 |
|  | medial | 4502 | 0 | 0.0 | 1672 | 0 | 0.0 | 2179 | 0 | 0.0 |
| **Total** |  | **40096** | **48** | **0.1** | **16688** | **15** | **0.1** | **22528** | **33** | **0.1** |

### The 5-HT2_C_ (Htr2c) receptor is expressed by both GABAergic and glutamatergic IC neurons

The 5-HT2_C_ receptor is an excitatory serotonergic receptor that is preferentially coupled to the G_q/11_ G-protein (Bockaert et al., 2006). These receptors are commonly located postsynaptically, although presynaptic expression has also been reported (Araneda and Andrade, 1991; Stanford and Lacey, 1996; Becamel et al., 2001). In the IC, activation of the 5-HT2_C_ receptor in vivo using selective agonists leads to an enhancement in firing rate in response to auditory stimuli (Hurley, 2006). Therefore, it has been hypothesized that the 5-HT2_C_ receptor is expressed by IC glutamatergic neurons (Hurley, 2006).

Here, we performed in situ hybridization using IC brain slices of three males and two females C57BL/6 mice and found that *Htr2c* is expressed by ∼46% (3748 out of 8093) of IC GABAergic neurons. (**Figure 5A-D, I, Table 9**). Additionally, our data show that ∼22.5% (13009 out of 57632) of glutamatergic neurons express *Htr2c* (**Figure 5E-H, J, Table 10**). The percentage of *Vgat^+^* neurons expressing *Htr2c* was higher across the whole IC as well as the central nucleus and shell, when compared to the percentage of *Vglut2*+ neurons expressing *Htr2c* (**Figure 5K**, IC: 46.2 ± 7.4% of *Vgat^+^Htr2c^+^ vs* 22.4 ± 5.3% of *Vglut^+^Htr2c^+^*, Welch’s *t test,* α = 3 x 10^-10^; ICc: 50.1 ± 11.7% of *Vgat^+^Htr2c^+^ vs* 14.7 ± 5.6% of *Vglut^+^Htr2c^+^*, Welch’s *t test,* α = 3 x 10^-10^; ICx: 43.1 ± 6.6% of *Vgat^+^Htr2c^+^ vs* 30.7 ± 9.8% of *Vglut^+^Htr2a^+^*, Welch’s *t test,* α = 0.001).

**Figure 5.**
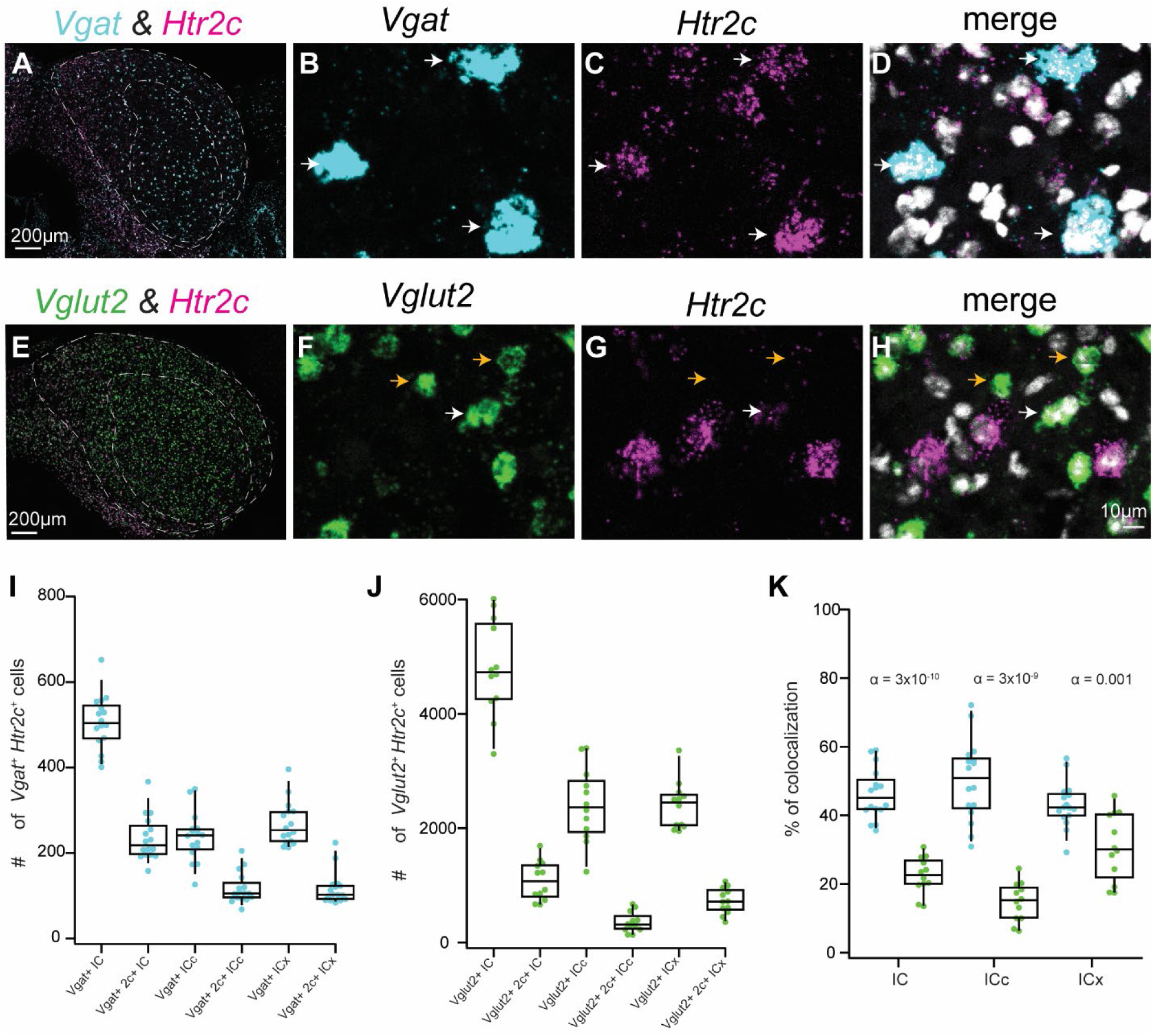
The 5-HT2C (Htr2c) receptor is expressed by both GABAergic and glutamatergic IC neurons. Fluorescent in situ hybridization was used to determine the expression patterns of Vgat, Vglut2, and Htr2c in the C57BL/6J mice. **A.** Coronal IC section showing the distribution of Vgat (cyan) and Htr2c (magenta) in the IC. **B-D.** High magnification confocal images showing that Vgat^+^ neurons highly colabel with Htr2c, as shown by white arrows. **E.** Coronal IC section showing the distribution of Vglut2 (green) and Htr2c (magenta) in the IC. **F-H.** High magnification confocal images showing that Vglut2^+^ neurons often do not colabel with Htr2a, as shown by orange arrows. However, some Vglut2^+^ neurons colabel with Htr2c, as shown by the white arrows. **I.** Boxplot showing the number of Vgat^+^ neurons and Vgat^+^ neurons that express the Htr2c in the IC, as well as in the central nucleus of the IC and shell IC. **J.** Boxplot showing the number of Vglut2^+^ neurons and Vglut2^+^ neurons that express the Htr2c in the IC, as well as in the central nucleus of the IC and shell IC. **K.** Comparison of the percentage of Vgat^+^ neurons expressing Htr2c (cyan dots) vs the percentage of Vglut2^+^ neurons expressing Htr2c (green dots) in the whole IC, central nucleus of the IC, and shell IC. α-values from Welch’s t-tests are atop each plot, showing the difference between the percentage of Vgat^+^ vs Vglut2^+^expressing Htr2c.

**Table 9.** Nearly half of IC GABAergic neurons express *Htr2c*.

| Mouse | Slice | *Vgat*+ | *Vgat*+ & *Htr2c*+ | % of co-label | *Vgat*+(ICc) | *Vgat*+ & *Htr2c*+(ICc) | % of co-label | *Vgat*+(ICx) | *Vgat*+ & *Htr2c*+(ICx) | % of co-label |
| --- | --- | --- | --- | --- | --- | --- | --- | --- | --- | --- |
| Male P45 | caudal | 529 | 227 | 42.9 | 218 | 104 | 47.7 | 311 | 123 | 39.5 |
|  | medial | 554 | 231 | 41.7 | 257 | 105 | 40.9 | 297 | 126 | 42.4 |
|  | medial | 427 | 158 | 37.0 | 202 | 68 | 33.7 | 225 | 90 | 40.0 |
| Male P45 | caudal | 414 | 196 | 47.3 | 174 | 97 | 55.7 | 240 | 99 | 41.3 |
|  | caudal | 509 | 245 | 48.1 | 212 | 117 | 55.2 | 297 | 128 | 43.1 |
|  | medial | 492 | 209 | 42.5 | 224 | 96 | 42.9 | 268 | 113 | 42.2 |
| Male P45 | caudal | 499 | 209 | 41.9 | 250 | 120 | 48.0 | 249 | 89 | 35.7 |
|  | rostral | 539 | 192 | 35.6 | 251 | 108 | 43.0 | 288 | 84 | 29.2 |
|  | rostral | 464 | 193 | 41.6 | 242 | 91 | 37.6 | 222 | 102 | 45.9 |
| Female P59 | caudal | 401 | 196 | 48.9 | 173 | 93 | 53.8 | 228 | 103 | 45.2 |
|  | medial | 563 | 294 | 52.2 | 350 | 205 | 58.6 | 213 | 89 | 41.8 |
|  | rostral | 557 | 207 | 37.2 | 343 | 106 | 30.9 | 214 | 101 | 47.2 |
| Female P59 | caudal | 526 | 255 | 48.5 | 283 | 163 | 57.6 | 243 | 92 | 37.9 |
|  | caudal | 498 | 294 | 59.0 | 240 | 173 | 72.1 | 258 | 121 | 46.9 |
|  | caudal | 652 | 367 | 56.3 | 256 | 143 | 55.9 | 396 | 224 | 56.6 |
|  | caudal | 469 | 275 | 58.6 | 126 | 87 | 69.0 | 343 | 188 | 54.8 |
| Total |  | 8093 | 3748 | 46.2 | 3801 | 1876 | 50.2 | 4292 | 1872 | 43.1 |

**Table 10.** Glutamatergic neurons expressing *Htr2c* are primarily in the shell IC.

| Mouse | Slice | *Vglut2*+ | *Vglut2*+ & *Htr2c*+ | % of co-label | *Vglut2*+(ICc) | *Vglut2*+ & *Htr2c*+(ICc) | % of co-label | *Vglut2*+(ICx) | *Vglut2*+ & *Htr2c*+(ICx) | % of co-label |
| --- | --- | --- | --- | --- | --- | --- | --- | --- | --- | --- |
| Male P45 | caudal | 6010 | 1691 | 28.1 | 3384 | 623 | 18.4 | 2627 | 1068 | 40.7 |
|  | caudal | 5900 | 1385 | 23.5 | 3399 | 674 | 19.8 | 2501 | 711 | 28.4 |
|  | medial | 4690 | 1445 | 30.8 | 2740 | 554 | 20.2 | 1950 | 891 | 45.7 |
| Female 59 | caudal | 3299 | 663 | 20.1 | 1240 | 304 | 24.5 | 2059 | 359 | 17.4 |
|  | caudal | 4224 | 858 | 20.3 | 2168 | 136 | 6.3 | 2056 | 722 | 35.1 |
|  | medial | 5498 | 744 | 13.5 | 2938 | 295 | 10.0 | 2560 | 449 | 17.5 |
| Female 59 | caudal | 4273 | 841 | 19.7 | 1767 | 231 | 13.1 | 2506 | 610 | 24.3 |
|  | caudal | 4767 | 669 | 14.0 | 1987 | 137 | 6.9 | 2780 | 532 | 19.1 |
|  | medial | 5674 | 1229 | 21.7 | 2314 | 232 | 10.0 | 3360 | 997 | 29.7 |
| Female P60 | caudal | 3827 | 923 | 24.1 | 1859 | 323 | 17.4 | 1968 | 600 | 30.5 |
|  | caudal | 4656 | 1222 | 26.2 | 2625 | 392 | 14.9 | 2031 | 830 | 40.9 |
|  | medial | 4814 | 1339 | 27.8 | 2418 | 377 | 15.6 | 2396 | 962 | 40.2 |
| Total |  | 57632 | 13009 | 22.5 | 28839 | 4278 | 14.8 | 28794 | 8731 | 30.8 |

GABAergic and glutamatergic neurons expressing *Htr2c* were present in all IC subdivisions. However, there was a small but statistically significant percentage of GABAergic neurons expressing *Htr2c* in the central nucleus of the IC compared to the shell IC (ICc: 50.1 ± 11.7% of *Vgat^+^Htr2c^+^ vs* ICx: 43.1 ± 6.6% of *Vgat^+^Htr2c^+^* neurons, Welch’s *t test,* P = 0.04). In contrast, glutamatergic neurons expressing *Htr2c* were more likely to be present in the shell IC (ICc: 14.7 ± 5.6% of *Vglut2^+^Htr2c^+^* neurons *vs* ICx: 30.7 ± 9.8% of *Vglut2^+^Htr2c^+^,* Welch’s *t test,* P = 0.0001).

### The 5-HT7 (Htr7) receptor is primarily expressed by IC GABAergic neurons

The 5-HT7 receptors are preferentially coupled with G_s_ protein receptors. These receptors are expressed by fusiform cells in the DCN (Tang and Trussell, 2015), and a previous study briefly mentioned that *Htr7* is also expressed in the IC (Heidmann et al., 1998). Here we found that ∼30% of IC GABAergic neurons *Htr7* (**Figure 6A-D, I, Table 11**). Next, we found that this receptor is also expressed by ∼12% of glutamatergic neurons (**Figure 6E-H, J, Table 12**). Our data show that the percentage of *Vgat^+^Htr7^+^* neurons was higher across the whole IC, central nucleus and shell IC, when compared to the percentage of *Vglut2*^+^ neurons expressing *Htr7* (**Figure 6K**, IC: 29.7 ± 8.3% of *Vgat^+^Htr7^+^ vs* 11.1 ± 5.9% of *Vglut^+^Htr7^+^*, Welch’s *t test,* α = 7 x 10^-5^; ICc: 28.2 ± 11.1% of *Vgat^+^Htr7^+^ vs* 9.2 ± 5.1% of *Vglut^+^Htr7^+^*, Welch’s *t test,* α = 0.006; ICx: 31.0 ± 10.0% of *Vgat^+^7^+^ vs* 14.2 ± 7.8% of *Vglut^+^Htr7^+^*, Welch’s *t test,* α = 0.001).

**Figure 6.**
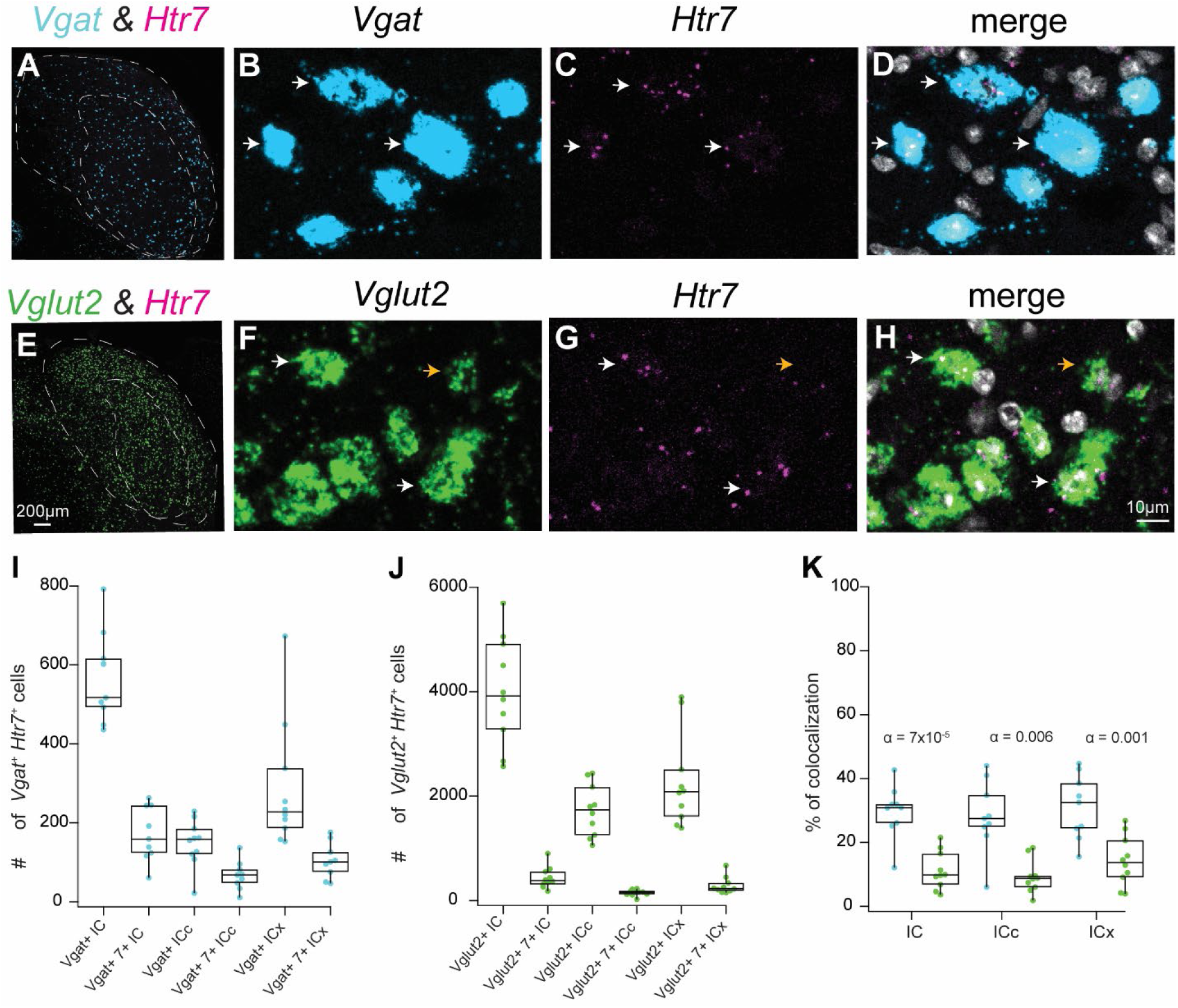
The 5-HT7 (Htr7) receptor is primarily expressed by IC GABAergic neurons. Fluorescent in situ hybridization was used to determine the expression patterns of Vgat, Vglut2, and Htr7 in the C57BL/6J mice. **A.** Coronal IC section showing the distribution of Vgat (cyan) and Htr7 (magenta) in the IC. **B-D.** High magnification confocal images showing that Vgat^+^ neurons often colabel with Htr7, as shown by white arrows. **E.** Coronal IC section showing the distribution of Vglut2 (green) and Htr7 (magenta) in the IC. **F-H.** High magnification confocal images showing that Vglut2^+^ neurons often do not colabel with Htr7, as shown by orange arrows. However, some Vglut2^+^ neurons colabel with Htr7, as shown by the white arrows. **I.** Boxplot showing the number of Vgat+ neurons and Vgat+ neurons that express the Htr7 in the IC, as well as in the central nucleus of the IC and shell IC. **J.** Boxplot showing the number of Vglut2^+^ neurons and Vglut2^+^ neurons that express the Htr7 in the IC, as well as in the central nucleus of the IC and shell IC. **K.** Comparison of the percentage of Vgat^+^ neurons expressing Htr7 (cyan dots) vs the percentage of Vglut2^+^ neurons expressing Htr7 (green dots) in the whole IC, central nucleus of the IC, and shell IC. α-values from Welch’s t-tests are atop each plot, showing the difference between the percentage of Vgat^+^ vs Vglut2^+^expressing Htr7.

**Figure 7.**
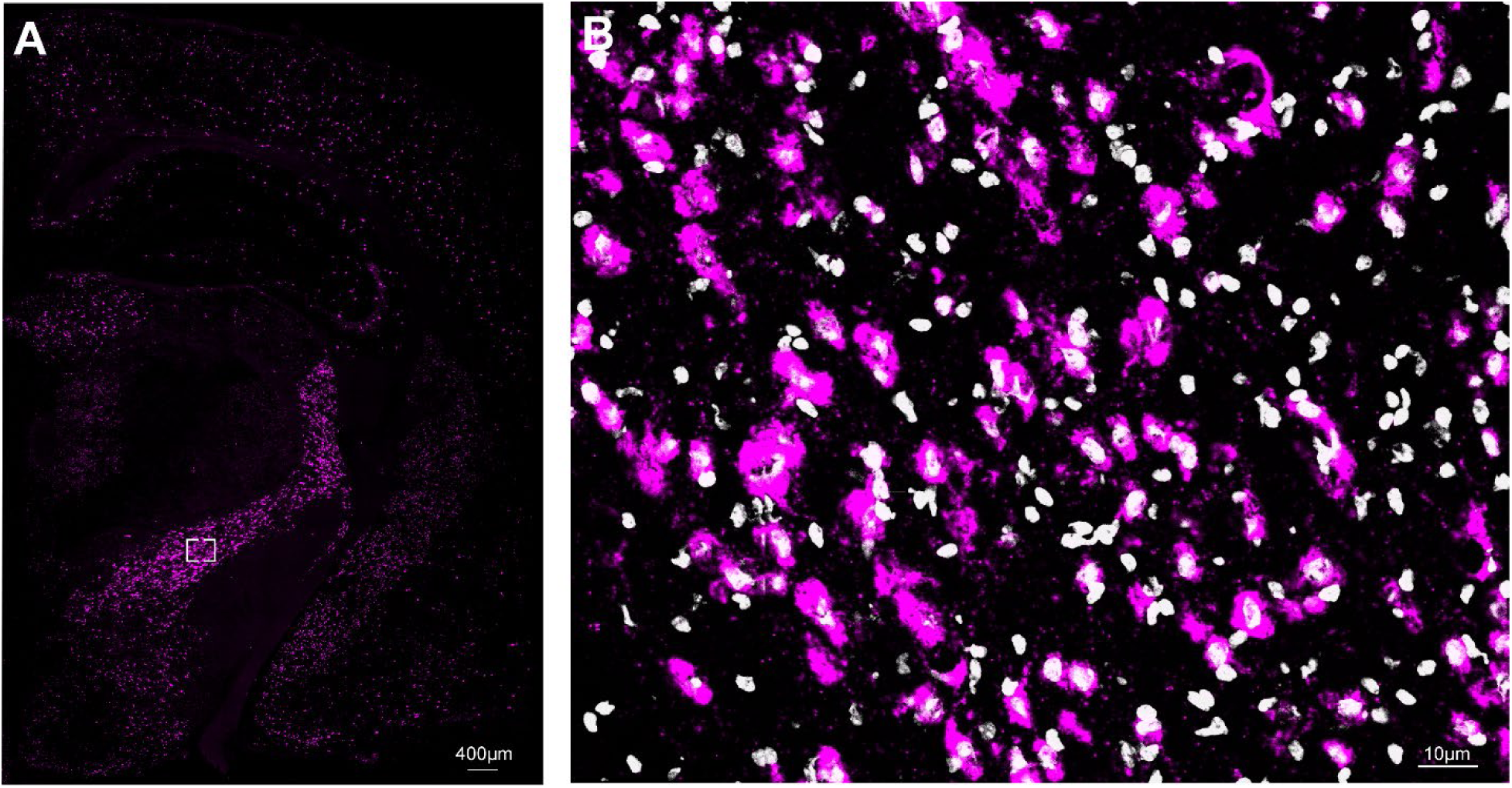
The 5-HT7 (Htr7) receptor is highly expressed in the cortex and hypothalamic regions. **A.** Confocal image showing that the Htr7 receptor is expressed in the cortex, zona incerta of the hypothalamus, and arcuate nucleus. The white rectangle represents the area of high magnification in **B.**

**Table 11.** Nearly 30% of IC GABAergic neurons express *Htr7*.

| Mouse | Slice | *Vgat*+ | *Vgat*+ & *Htr7* | % of co-label | *Vgat*+(ICc) | *Vgat*+ & *Htr7*+(ICc) | % of co-label | *Vgat*+(ICx) | *Vgat*+ & *Htr7*+(ICx) | % of co-label |
| --- | --- | --- | --- | --- | --- | --- | --- | --- | --- | --- |
| **Male P50** | caudal | 682 | 244 | 35.8 | 273 | 68 | 24.9 | 409 | 176 | 43.0 |
|  | medial | 436 | 139 | 31.9 | 132 | 34 | 25.8 | 304 | 105 | 34.5 |
|  | medial | 792 | 245 | 30.9 | 369 | 82 | 22.2 | 423 | 163 | 38.5 |
|  | rostral | 616 | 263 | 42.7 | 334 | 137 | 41.0 | 282 | 126 | 44.7 |
| **Male P50** | medial | 602 | 192 | 31.9 | 218 | 96 | 44.0 | 384 | 96 | 25.0 |
|  | rostral | 517 | 159 | 30.8 | 206 | 58 | 28.2 | 311 | 101 | 32.5 |
| **Female 45** | caudal | 448 | 117 | 26.1 | 255 | 70 | 27.5 | 193 | 47 | 24.4 |
|  | medial | 506 | 61 | 12.1 | 183 | 11 | 6.0 | 323 | 50 | 15.5 |
|  | rostral | 493 | 124 | 25.2 | 138 | 48 | 34.8 | 355 | 76 | 21.4 |
| **Total** |  | **5092** | **1544** | **29.7** | **2108** | **604** | **28.3** | **2984** | **940** | **31.1** |

**Table 12.** Nearly 12% of IC glutamatergic neurons express *Htr7*.

| Mouse | Slice | *Vglut2*+ | *Vglut2*+ & *Htr7*+ | % of co-label | *Vglut2*+(ICc) | *Vglut2*+ & *Htr7*+(ICc) | % of co-label | *Vglut2*+(ICx) | *Vglut2*+ & *Htr7*+(ICx) | % of co-label |
| --- | --- | --- | --- | --- | --- | --- | --- | --- | --- | --- |
| **Male P50** | caudal | 2668 | 439 | 16.5 | 1059 | 185 | 17.5 | 1609 | 254 | 15.8 |
|  | medial | 3276 | 371 | 11.3 | 1834 | 162 | 8.8 | 1442 | 209 | 14.5 |
|  | medial | 4915 | 902 | 18.4 | 2407 | 229 | 9.5 | 2508 | 673 | 26.8 |
|  | rostral | 2571 | 554 | 21.5 | 1182 | 216 | 18.3 | 1389 | 338 | 24.3 |
| **Male P50** | medial | 3851 | 605 | 15.7 | 1672 | 156 | 9.3 | 2179 | 449 | 20.6 |
|  | medial | 3987 | 395 | 9.9 | 2175 | 161 | 7.4 | 1812 | 234 | 12.9 |
|  | rostral | 5055 | 180 | 3.6 | 1256 | 22 | 1.8 | 3799 | 158 | 4.2 |
| **Female P45** | caudal | 3577 | 348 | 9.7 | 1477 | 127 | 8.6 | 2100 | 221 | 10.5 |
|  | caudal | 5694 | 261 | 4.6 | 1798 | 108 | 6.0 | 3896 | 153 | 3.9 |
|  | medial | 4502 | 308 | 6.8 | 2436 | 121 | 5.0 | 2066 | 187 | 9.1 |
| **Total** |  | **40096** | **4363** | **11.8** | **17296** | **1487** | **9.2** | **22800** | **2876** | **14.3** |

Across all IC subdivisions, GABAergic and glutamatergic neurons were found to express *Htr7*. The percentage of GABAergic neurons expressing *Htr7* in the central nucleus of the IC was not statistically different when compared to the shell IC (ICc: 28.2 ± 11.1% of *Vgat^+^Htr7^+^ vs* ICx: 31.0 ± 10.0% of *Vgat^+^Htr7^+^* neurons, Welch’s *t test,* P = 0.58). Glutamatergic neurons expressing *Htr7* were also equally distributed across the central nucleus of the IC and shell IC (ICc: 9.2 ± 5.1% of *Vglut2^+^Htr7^+^* neurons *vs* ICx: 14.2 ± 7.8% of *Vglut2^+^Htr7^+^,* Welch’s *t test,* P = 0.10).

Although the *Htr7* was expressed in the IC, the qualitative expression was lower compared to the other receptors shown here. Additionally, many cells expressed only 3-4 puncta, which was the lowest threshold for us to consider a cell positive for a specific receptor. This data may suggest that although *Htr7* is present in the IC, the level of expression is lower compared to other serotonergic receptors. To ensure that our reaction was working correctly, we included slices containing cortex, hippocampus, and hypothalamic regions in our assay, in which 5-HT7 has been well validated (Hedlund and Sutcliffe, 2004; Siddiqui et al., 2004; Roberts and Hedlund, 2012a). As shown in **Figure 7**, we found that *Htr7* is highly expressed in the cortex and hypothalamic regions, including the arcuate nuclei and zona incerta of the hypothalamus.

### Colocalization of serotonergic receptors in the IC

Co-expression of serotonergic receptors has been reported in many brain regions, including the dorsal cochlear nucleus, where fusiform cells co-express 5-HT2_A_, 5-HT2_C_, and 5-HT7 (Tang and Trussell, 2015). In addition, more complex co-expression of serotonergic receptors has been shown in pyramidal neurons in the prefrontal cortex, which co-express 5-HT1_A_ and 5-HT2_A,_ suggesting an excitatory and inhibitory effect of serotonin on the same cell (Amargós-Bosch et al., 2004).

To determine whether serotonergic receptors are co-expressed in the IC, we used different combinations of probes in our in-situ hybridization assays: (1) *Vgat*, *Htr1b*, and *Htr2a*, (2) *Vgat*, *Htr1a*, and *Htr2c*, (3) *Vgat*, *Htr1a*, and *Htr1b*, (4) *Vgat*, *Htr2a*, and *Htr2c*, (5) *Vglut2*, *Htr1a*, and *Htr1b* and (6) *Vglut2*, *Htr2a*, and *Htr2c.* We found that inhibitory and excitatory serotonin receptors are rarely co-expressed in single IC neurons. Out of 6402 *Vgat^+^* neurons, only 72 (1.2%) co-express *Htr1b* and *Htr2a.* In agreement, less than 2% (79 out of 4427) of *Vgat^+^* co-express *Htr1a* and *Htr2c*. On the other hand, *Htr2a* and *Htr2c* were co-expressed by ∼30% of *Vgat^+^* neurons (1134 out of 3637, **Figure 8A-H**). Even though some glutamatergic neurons express *Htr2c,* the co-expression with *Htr2a* was rare (739 out of 57623, 1.3%).

**Figure 8.**
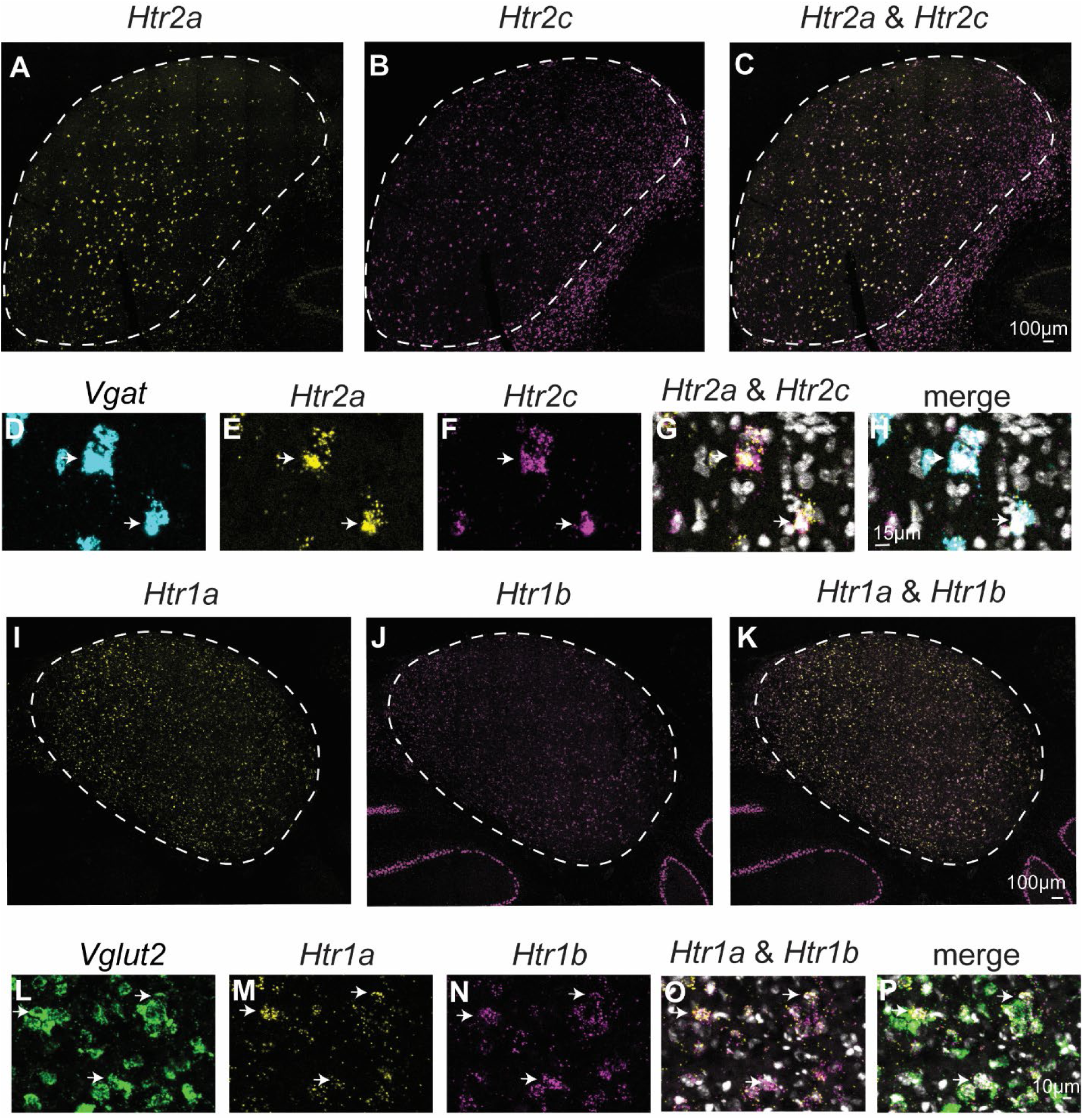
Colocalization of serotonergic receptors in the IC. A-C. Distribution of Htr2a (yellow) and Htr2c (magenta) in the IC. Merge in **C.** High magnification confocal images showing that Vgat+ neurons (**D**, cyan) often co-express Htr2a (**E**, yellow) and Htr2c (**F**, magenta). Merge in **G** and **H. I-K.** Distribution of Htr1a (yellow) and Htr1b (magenta) in the IC. Merge in **K.** High magnification confocal images showing that Vglut2+ neurons (**L**, green) often co-express Htr1a (**M**, yellow) and Htr1b (**N**, magenta). Merge in **O** and **P.**

Additionally, the inhibitory serotonergic receptors, *Htr1a* and *Htr1b*, were found to be co-expressed by ∼12% (9542 out of 78046) of *Vglut2^+^* cells (**Figure 8I-P**).

## Summary of results

Figure 9 summarizes the results of our in-situ hybridization studies. Together, our experiments demonstrate that four subtypes of metabotropic serotonergic receptors are highly expressed in the IC: *Htr1a, Htr1b, Htr2a,* and *Htr2c.* Those receptors were differentially expressed among GABAergic and glutamatergic neurons. GABAergic neurons primarily express the excitatory serotonergic receptors *Htr2a* and *Htr2c,* but glutamatergic neurons primarily express the excitatory serotonergic receptors *Htr1a* and *Htr1b.* These data suggest that serotonin activation of its receptors in the IC may lead to an inhibitory net effect.

**Figure 9.**
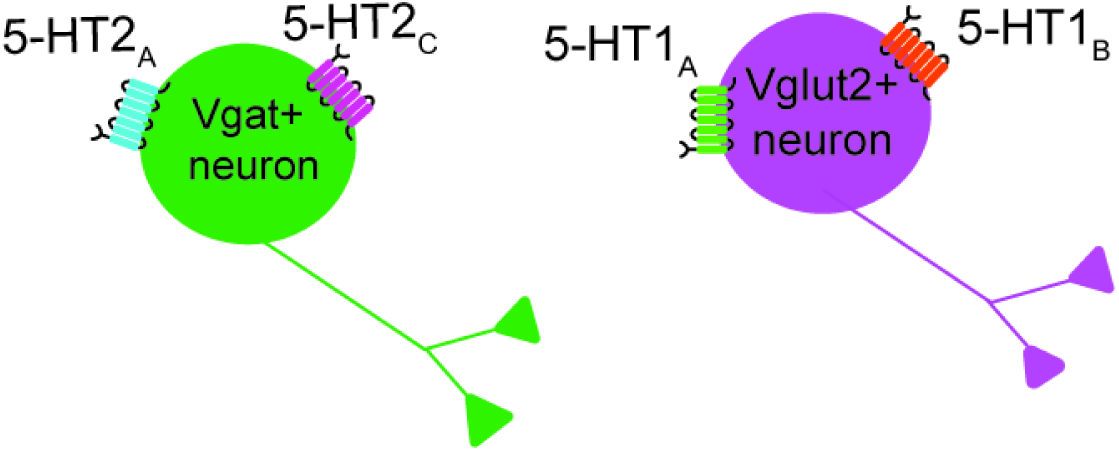
Summary of results. Together, our results suggest that GABAergic neurons (Vgat^+^) primarily express excitatory serotonergic receptors while glutamatergic neurons (Vglut2^+^) primarily express inhibitory serotonergic receptors.

## Discussion

Here, we showed that the metabotropic serotonergic receptors *Htr1a, Htr1b, Htr2a,* and *Htr2c* are widely distributed throughout the IC. Interestingly, we see less expression of *Htr7* and nearly no expression of *Htr2b.* We next found that *Htr1a* and *Htr1b,* which are inhibitory serotonergic receptors, largely co-label with glutamatergic IC neurons. However, the expression of *Htr1a* and *Htr1b* was rarely observed in GABAergic neurons, identified by the expression of *Vgat.* On the other hand, the excitatory serotonergic receptor *Htr2a* was highly expressed by GABAergic neurons, but very few glutamatergic neurons were found to express this receptor.

Finally, *Htr2c,* which is also an excitatory serotonergic receptor, was primarily expressed by GABAergic neurons but also expressed by many glutamatergic neurons. Together, our data provide evidence regarding the distribution of six subtypes of serotonergic receptors across GABAergic and glutamatergic neurons in the IC. Since serotonergic neurons are concentrated in the DRN, the net effect of serotonin on IC sound processing will be defined by the interaction of different receptors expressed by neuronal classes. Given the differential expression of those receptors by GABAergic and glutamatergic neurons in the IC, serotonin is likely to have an inhibitory net effect.

### IC glutamatergic neurons primarily express Htr1a and Htr1b

5-HT1_A_ receptors play a major role as autoreceptors in the DRN, regulating the firing of serotonergic neurons (Riad et al., 2000; Andrade et al., 2015). Outside of the DRN, 5-HT1_A_ receptors are primarily postsynaptic and, when activated by serotonin, inhibit neuronal activity through activation of GIRK channels and/or the inhibition of calcium channels (Andrade et al., 1986; Okuhara and Beck, 1994). Autoradiographic studies suggested that the IC contains the highest number of 5-HT1_A_ binding sites among the central auditory pathway (Thompson et al., 1994b). Previous studies using in vivo electrophysiology have shown that the application of the 5-HT1_A_ agonist decreases evoked spike rates and narrows the frequency tuning of IC neurons (Hurley, 2006; Ramsey et al., 2010). This data is consistent with our findings that IC glutamatergic neurons primarily express *Htr1a*, and activation of 5-HT1_A_ receptors would likely hyperpolarize these glutamatergic neurons, thereby decreasing excitability in the IC. However, another study used antibodies to identify 5-HT1_A_ and found that ∼50% of IC GABAergic neurons co-label with 5-HT1_A_ (Peruzzi and Dut, 2004). This discrepancy among results is likely due to the difference in techniques used, since antibodies can be less specific due to the high similarity of serotonergic receptors.

In contrast to 5-HT1_A_ receptors, 5-HT1_B_ receptors are more likely to be expressed in neuron terminals, and activation of the 5-HT1_B_ receptors inhibits neurotransmitter release. The 5-HT1_B_ receptor has been shown to inhibit the release of dopamine, GABA, and glutamate in many brain regions, including the striatum and hippocampus (Pommer et al., 2021; Burke and Alvarez, 2022; Najm Al-Halboosi et al., 2023). In the IC, it has been suggested that GABAergic neurons express 5-HT1_B_. This hypothesis is supported by immunohistochemistry experiments as well as in vivo electrophysiology using the 5-HT1_B_ agonist and/or antagonist (Peruzzi and Dut, 2004; Hurley et al., 2008). Application of the 5-HT1_B_ agonist reduces the number of spikes in a frequency-specific manner, and this effect is diminished in the presence of the GABA_A_ receptor antagonist (Hurley et al., 2008). This contradicts the data from the current study since we observed minimal expression of *Htr1b* in *Vgat+* neurons (Figure 2). A possible explanation for this discrepancy may be that glutamatergic neurons are largely interconnected with other IC neurons (Sturm et al., 2014, 2017; Ito et al., 2016; Oberle et al., 2023; Silveira et al., 2023).

Thereby, serotonin modulation of glutamate transmission could indirectly affect the membrane excitability of IC GABAergic neurons, bringing these neurons under threshold, reducing the release of GABA. In agreement with the data presented here, whole-brain gene expression analysis has found that indeed, few IC GABAergic neurons express *Htr1b* (Lein et al., 2007). However, those are highly expressed by GABAergic Purkinje cells in the cerebellum (Lein et al., 2007). As shown in Figure 1B, our in-situ hybridization can detect high expression of H*tr1b* in GABAergic Purkinje cells, thereby supporting the low colocalization of H*tr1b* with GABAergic cells in the IC.

### Over half of IC GABAergic neurons express Htr2a

The 5-HT2_A_ receptors have received much attention due to their involvement in hallucinations and schizophrenia (Michaiel et al., 2019; Nakao et al., 2022). In the central auditory system, 5-HT2_A_ receptors have been shown to modulate the excitability of fusiform cells in the dorsal cochlear nucleus (Tang and Trussell, 2015, 2017). A previous study also demonstrated that the application of the 5-HT2_A_ agonist increases the frequency of inhibitory postsynaptic currents (IPSCs) in the IC (Wang et al., 2008). This finding aligns with our data, which shows that over half of IC GABAergic cells co-express *Htr2a.* Expression of *Htr2a* was rarely observed in glutamatergic neurons. When present, it had a lower intensity and a decreased number of puncta compared to its expression in GABAergic neurons (data not shown). Although more functional data is necessary, this suggests a significant role of 5-HT2_A_ in regulating local inhibition in the IC and raises interesting hypotheses, such as the involvement of this receptor in auditory hallucinations.

### Both GABAergic and glutamatergic neurons express Htr2c

The 5-HT2_C_ receptor is an excitatory serotonergic receptor, and it is mainly expressed postsynaptically, but has also been shown to be present presynaptically (Araneda and Andrade, 1991; Stanford and Lacey, 1996). For example, in the striatum, activation of the 5-HT2_C_ receptor by serotonin produces an inhibitory effect since this receptor is expressed by fast-spiking GABAergic interneurons (Blomeley and Bracci, 2009). In the IC, *Htr2c* is upregulated after noise exposure (Holt et al., 2005). In addition, application of the 5-HT2_C_ agonist in vivo using juxtacellullar recordings induces an increase in firing rate. However, those recordings were not selectively targeted to either GABAergic or glutamatergic neurons (Hurley, 2006). In our data, we found that ∼20% of IC glutamatergic neurons express *Htr2c.* However, this receptor is also expressed by ∼50% of GABAergic neurons. Thereby, the activation of this receptor by serotonin is likely to have a more complex net effect since it will affect both glutamatergic and GABAergic neurons. A possible explanation for the discrepancy between our data and previous studies is that the majority of neurons in the IC are glutamatergic. Therefore, in vivo recordings are more likely to target IC glutamatergic neurons.

### 5-HT7 receptors are primarily expressed by GABAergic neurons

The 5-HT7 is an excitatory metabotropic serotonergic receptor that is preferentially coupled with G_s_ protein receptors. Activation of these receptors leads to an enhancement in neuronal excitability. A physiological role for the 5-HT7 receptor has been shown in circadian rhythm regulation (Shelton et al., 2015), thermoregulation (Voronova, 2021), and learning and memory (Roberts and Hedlund, 2012b). In the central auditory pathway, activation of 5-HT7 receptors enhances the excitability of fusiform cells in the DCN. In addition, a previous study briefly mentioned that *Htr7* is expressed in the IC (Heidmann et al., 1998). Here we found that ∼30% of GABAergic neurons express the *Htr7,* while ∼12% of glutamatergic neurons express this receptor. Although *Htr7* expression was present in the IC, we found it to have a lower qualitative expression than the other receptors reported here. In many cases, a positive cell had only a minimal number of puncta (3-4) to be considered positive. However, quantitative puncta analysis will be necessary, using samples that include other serotonergic receptors in the same assay, to confirm this hypothesis.

### Distribution of serotonergic receptors across the central nucleus of the IC and the shell of the IC

The IC is subdivided into three main subdivisions: the central nucleus (ICc), dorsal cortex (ICd), and lateral cortex (ICl). In this study, we analyzed the distribution of serotonergic receptors across the central and shell IC (ICd + ICl). We found that all serotonergic receptors are present throughout the IC, suggesting that serotonergic modulation acts on lemniscal and non-lemniscal circuits. Some receptors, such as *Htr1b*, showed stronger expression in the shell IC compared to the central IC. Interestingly, *Htr2c* was preferentially expressed by GABAergic neurons in the central IC, but when present in glutamatergic neurons, it was more often localized to the shell IC. These data suggest that *Htr2c* may play an important role in integrating multisensory inputs by modulating glutamatergic neurons in the shell IC, but likely influences sound processing in the central IC through the modulation of GABAergic neurons. A limitation of these findings is that different approaches to identifying the IC subdivisions can lead to different limits between the central and shell that could impact the interpretation of the data. Here, we used the GAD67 and Glyt2 immunohistochemistry staining to identify the subdivisions. Different groups have validated this method (Choy Buentello et al., 2015; Beebe et al., 2020; Silveira et al., 2020). However, other approaches to identifying the IC subdivisions have also been previously reported and could influence how to define the limits of the IC borders (Keesom et al., 2018; Olthof et al., 2019).

### Functional implications: a possible role for serotonin in regulating E/I balance

The serotonergic system has been shown to regulate EI balance in other brain regions (Moreau et al., 2010; Carlos-Lima et al., 2023). Here, we show that in the IC, metabotropic serotonergic receptors are differentially distributed across GABAergic and glutamatergic neurons. *Htr1a* and *Htr1b* (inhibitory serotonergic receptors) were primarily expressed by glutamatergic neurons, and *Htr2a* (excitatory serotonergic receptor) was almost exclusively expressed by GABAergic neurons. *Htr2c* was expressed by both GABAergic and glutamatergic neurons, although glutamatergic neurons expressing *Htr2c* were primarily in the shell IC. Together, these data suggest that serotonin modulation induces an inhibitory net effect in the IC by differentially affecting glutamatergic and GABAergic neurons. This proposed mechanism may help to understand how some auditory disorders lead to hyperexcitability in the IC. For example, a recent study showed that activation of the 5-HT1_A_ receptor in the IC prevents audiogenic seizures in the Fmr1 knockout mice, a mouse line used to study fragile X syndrome (Saraf et al., 2024). Based on our findings, this effect would likely be achieved by hyperpolarization of IC glutamatergic neurons, which are necessary for the initiation of audiogenic seizures (Gonzalez et al., 2019). In addition, dysfunction in serotonergic modulation is involved in the perception and generation of tinnitus (Salvinelli et al., 2003; Caperton and Thompson, 2010). Furthermore, medications that selectively inhibit the reuptake of serotonin (SSRIs), commonly used in the treatment of depression (Stark et al., 1985), have been shown to improve tinnitus perception (Shemen, 1998). However, SSRIs affect all serotonergic transmission and can cause multiple side effects (Vaswani et al., 2003). Therefore, the ability to identify which subtypes of receptors are expressed by inhibitory and excitatory neurons may lead to the development of more targeted therapeutics. For example, it was recently shown that modulation of the 5-HT1_A_ receptor in the auditory cortex improves temporal processing in a mouse model of autism (Tao et al., 2025). Many central auditory disorders are characterized by changes in excitability that favor excitation, such as tinnitus, age-related hearing loss, and audiogenic seizure (Auerbach et al., 2014; Xiong et al., 2017; Gonzalez et al., 2019). Since our data show that serotonin favors inhibition in the IC by differentially regulating GABAergic and glutamatergic neurons, dysfunction in serotonergic modulation could potentially underlie enhanced excitability in central auditory disorders.

## Conclusions

We demonstrate that four subtypes of metabotropic serotonergic receptors are highly expressed in the IC: *Htr1a, Htr1b, Htr2a,* and *Htr2c.*GABAergic neurons were more likely to express the excitatory serotonergic receptors *Htr2a* and *Htr2c,* but glutamatergic neurons primarily express the excitatory serotonergic receptors *Htr1a* and *Htr1b,* although the *Htr2c* receptor was also present in glutamatergic neurons. These data suggest that by differentially regulating GABAergic and glutamatergic neurons, serotonin activation of its receptors may lead to an inhibitory net effect.

## Author statement

Marina Silveira: Conceptualization, Investigation, Writing - Original draft, Writing - Review & editing, Visualization, Supervision, Funding Acquisition; Karen Galindo: Investigation, Writing - Review & editing; Zoya Nazir: Investigation, Writing - Review & editing.

## Notes

**Acknowledgements:** This work was supported by National Institutes of Health Grants NIH R00 DC019415 (MAS), UTSA RISE Program NIH/NIGMS GM060655 (KLG), and UTSA funding. We thank all the members of the Silveira Lab for their feedback. We also thank Elie Huez and Dr. Audrey Drotos for comments on the manuscript.

### Competing Interest Statement

The authors have declared no competing interest.

